# GmVTL1 is an iron transporter on the symbiosome membrane of soybean with an important role in nitrogen fixation

**DOI:** 10.1101/2020.03.03.975805

**Authors:** Ella M. Brear, Frank Bedon, Aleksandr Gavrin, Igor S. Kryvoruchko, Ivone Torres-Jerez, Michael K. Udvardi, David A. Day, Penelope M.C. Smith

**Author notes:** Corresponding author: Penelope M. C. Smith.

## Abstract

- Legumes establish symbiotic relationships with soil bacteria (rhizobia), housed in nodules on plant roots. The plant supplies carbon substrates and other nutrients to the bacteria in exchange for fixed nitrogen. The exchange occurs across a plant-derived symbiosome membrane (SM), which encloses rhizobia to form a symbiosome. Iron supplied by the plant is crucial for the rhizobial enzyme nitrogenase that catalyses N_2_ fixation, but the SM iron transporter has not been identified.
- We use complementation of yeast and plant mutants, real-time PCR, hairy root transformation, microscopy and proteomics to demonstrate the role of soybean GmVTL1 and 2.
- Both are members of the vacuolar iron transporter family and homologous to *Lotus japonicus* SEN1 (LjSEN1), previously shown to be essential for N_2_ fixation. *GmVTL1* expression is enhanced in nodule infected cells and both proteins are localised to the SM.
- GmVTL1 and 2 transport iron in yeast and GmVTL1 restores N_2_ fixation when expressed in the *Ljsen1* mutant.
- Three GmVTL1 amino acid substitutions that reduce iron transport in yeast also block N_2_ fixation in *Ljsen1* plants.
- We conclude GmVTL1 is responsible for transport of iron across the SM to bacteroids and plays a crucial role in the N_2_-fixing symbiosis.

## Introduction

Iron is an essential plant micronutrient, important for both respiration and photosynthesis. In cells it is often complexed with other molecules and sequestered in organelles to prevent accumulation of toxic levels of the free form in the cytosol. One strategy used to constrain iron concentrations in the cytosol is to sequester excess in the vacuole. Members of the CCC/Vacuolar iron transporter (VIT) family are responsible for detoxification of iron by transporting it into the vacuole in yeast, plants and *Plasmodium falciparum* (Li *et al.*, 2001, Kim *et al*. 2006, Labarbuta *et al*., 2017). The mechanism of transport by the VIT proteins of *P. falciparum* and *Eucalyptus grandis* appears to be proton coupled antiport, taking iron into the vacuole in exchange for a proton (Labarbuta *et al*., 2017, Kato et al. 2019). One of the first members of this gene family to be described was Nodulin 21, which was expressed specifically in nodules of soybean that had formed a symbiosis with nitrogen-fixing rhizobia (Delauney *et al.*, 1990). More recently, a mutant for another VIT family member, *sen1* (*<u>s</u>*tationary *<u>e</u>*ndosymbiont *<u>n</u>*odule), was described in *Lotus japonicus*. Mutations in *Ljsen1* block nitrogen fixation, suggesting an important role for this gene family in the legume:rhizobia symbiosis (Suganuma *et al*., 2003; Hakoyama *et al.*, 2012). However, although the effects of the mutation in *sen1* are well documented, the subcellular localisation and function of the LjSEN1 protein have not been determined.

In the symbiosis, N_2_ fixed by the rhizobial enzyme, nitrogenase, is exchanged for supply of a carbon source (probably malate) and all the nutrients required for nitrogen fixation and bacterial metabolism (Udvardi & Day, 1997). This exchange occurs in a new organ, the nodule, that forms to house the rhizobia. Within this organ, rhizobia differentiate into their symbiotic form, bacteroids, and are compartmentalized from the plant cytoplasm by a plant-derived membrane to form an organelle-like structure, the symbiosome (Mohd-Noor et al., 2015). The plant membrane surrounding the bacteroids is known as the symbiosome membrane (SM).

Iron is essential for the symbiosis. It is required for both the iron- and molybdenum-iron proteins of the enzyme nitrogenase and for enzymes of the bacterial respiratory chain. Leghaemoglobin, with which the plant maintains the low free O_2_ concentration in infected cells required for nitrogenase activity, also contains iron (Guerinot, 1991; Brear 2013; González-Guerrero *et al.*, 2014). At maturity, the nodule contains the highest proportion of iron of all plant organs (Burton *et al.*, 1998). Iron taken up by the roots must be transported to the nodule and then directed to the cells and organelles that require it. Before it is available to the bacteroid, iron must first reach and be imported into infected cells and then transverse the SM and the space between the SM and the bacteroid membrane (Brear *et al*., 2013).

Most iron is imported into the nodule via the xylem (Rodriguez-Haas *et al*., 2013) and most likely as ferric citrate (Cline *et al*., 1982; Brear et al. 2013). Iron must move across a number of cell layers to reach the infected cells (Guinel, 2009) and both symplastic and apoplastic routes are utilised (Brown *et al*., 1995; Bederska *et al.*, 2012; Rodriguez-Haas *et al*., 2013). Ferrous iron uptake by the infected cell is facilitated by a member of the NRAMP family in *Medicago truncatula*, MtNRAMP1 (Tejada-Jiménez *et al*., 2015), possibly acting in concert with Multidrug And Toxic Compound Extrusion protein 67 (MtMATE67), which catalyses citrate efflux and appears to enhance iron uptake by infected cells (Kryvoruchko *et al*., 2018).

In soybean, both ferrous and ferric iron are transported across the SM in isolated symbiosomes (Moreau et al., 1995; LeVier et al., 1996; Moreau et al., 1998). The identification of ferric chelate reductase activity on the SM suggests ferrous iron is preferentially transported (LeVier et al., 1996). The symbiosome space in soybeans has high levels of non-heme iron bound to chromophores (Wittenberg *et al*., 1996) and may act as an iron store (Udvardi & Day, 1997). GmDMT1 (*Glycine max* Divalent Metal Transporter 1), a member of the NRAMP family, is localised on the SM and partially complemented a yeast mutant, *fet3fet4*, deficient in iron uptake (Kaiser *et al*., 2003). Although the direction it transports iron has not been determined *in vivo*, the fact it transports iron into the cytoplasm when expressed in yeast, together with the known characteristics of other NRAMP family transporters, suggest it is unlikely to catalyze uptake of iron into the symbiosome (equivalent to export from the cytoplasm) (Brear *et al.*, 2013; González-Guerrero *et al*., 2014). Rather, based on its phylogenetic similarity to *Arabidopsis thaliana* NRAMP3 and 4 that are localised on the tonoplast and remobilise stored iron during germination (Lanquar *et al*., 2004; Lanquar *et al*., 2005), and the fact that symbiosomes effectively replace the vacuole in infected cells (Whitehead and Day, 1997; Gavrin *et al*., 2014), GmDMT1 is more likely to transport iron from the symbiosome store to the plant cytoplasm (Brear *et al.*, 2013; González-Guerrero *et al*., 2014).

Since other members of the VIT family export iron out of the cytoplasm into vacuoles (Kim et al. 2006; Gollhofer et al. 2014; Connorton et al., 2017), Nodulin-21 and SEN1 are more likely candidates for iron uptake into the symbiosome than GmDMT1. Transcriptome data for soybean (Libault *et al*., 2010; Severin *et al*., 2010; Cao, 2019) identified two members of the VIT family with high expression in nodules (Brear *et al.*, 2013). One of these, Glyma.05G121600, corresponds to Nodulin 21 (Delauney *et al*., 1990). The second, Glyma.08G076300, encodes a protein with 87.8% identity to Glyma.05G121600. We have named these genes *GmVTL1* and *GmVTL2*, respectively.

Here we show that the soybean homologues of LjSEN1, GmVTL1 and GmVTL2 are able to transport iron into vacuoles when expressed in yeast and are localised to the SM in soybean nodules. GmVTL1 complements the *Ljsen1* mutant and mutation of conserved amino acids that in SEN1 block nitrogen fixation, reduce or eliminate iron transport by GmVTL1 in yeast. We propose that GmVTL1, and to a lesser extent GmVTL2, play a crucial role in transporting iron into the symbiosome to support nitrogen-fixing bacteroids.

## Materials and Methods

### Plant growth and hairy root transformation

*G. max* cv. Stephens seeds were inoculated with *B. japonicum* (Soybean group H, New Edge Microbials) and grown as described in Clarke et al. (2015). *L. japonicus sen1-1* (Suganuma *et al*., 2003; Hakoyama *et al.*, 2012) and Gifu plants were grown in 1:1 vermiculite: perlite and inoculated with *Mesorhizobium loti* strain NZP2235. Plants were fertilized with B & D nutrient solution (Broughton & Dilworth, 1971) without a nitrogen source.

The *Agrobacterium rhizogenes* hairy root transformation methods for *G. max* (cvs. Moonbi or Stephens) were described by Kereszt et al. (2007) and Mohammadi-Dehcheshmeh et al. (2014) and for *L. japonicus* by Díaz et al. (2005). Transgenic roots were visualized using a Leica M205FA stereomicroscope (Leica Microsystems) by detection of the DsRed fluorophore (excitation, 540/580; emission, 593/667).

### Cloning

The *GmVTL1* promoter was amplified from soybean genomic DNA, using two-step attB adaptor PCR (Life Technologies) with Phusion High Fidelity DNA Polymerase (NEB). Coding sequences of *GmVTL1* and *GmVTL2* with or without stop codon were amplified from nodule cDNA using Pfx50™ (Invitrogen). The coding sequence of *LjSEN1* was amplified from nodule cDNA using DNA polymerase KOD (EMD Millipore). Primers are listed in Supporting Information Table S1. Mutated *GmVTL1* coding regions were synthesized by Synbio Technologies.

Constructs were prepared using the Gateway cloning system (Invitrogen). The vector pKGW_GGRR_C was used for promoter GUS fusion (Ivanov *et al.*, 2012) and pGmlbc3-pK7WGF2-R (Gavrin *et al.*, 2016) for N-terminal GFP fusions. For yeast expression coding regions with or without stop were recombined into pDR196GW, pAG426GAL_eGFP_ccdB (Addgene; N-terminal GFP fusions) or pAG426GAL_ccdB_eGFP (Addgene; C-terminal GFP fusion).

### Quantitative real time PCR analysis

Total RNA was extracted from plant tissue using the RNeasy Plant mini kit (QIAGEN), with on-column DNase treatment. Complementary DNA (cDNA) was synthesized using the iScript cDNA synthesis kit (Bio-Rad) with primers (Supporting Information Table S1) designed with Primer3 (Untergasser *et al*., 2012). Real-time PCR reactions comprised 1 X SYBR-Green I master mix, 0.5 µM of each primer and 1 µl cDNA in a 5 µl reaction. Relative quantification analysis was performed using the LightCycler® 480 Real-Time PCR System software (v1.5.0 SP3, Roche Applied Science). Efficiency values for each amplicon were calculated by LinReg PCR software (Ramakers *et al*., 2003) and expression normalized against *GmUbi3* (Glyma.20G141600, Trevaskis *et al*., 2002).

### GUS staining, sample preparation and light microscopy

Transgenic nodules (21 DAI) were assessed for GUS staining as described in Clarke et al. (2015).

### Localisation using particle bombardment in leek

Mature leek plants (*Allium porrum* L.) were transiently transformed by particle bombardment following the method outlined in Collings *et al.* (2002), using 1 µm diameter gold particles (Bio-Rad Laboratories, USA) coated with 500 ng of each plasmid DNA via CaCl_2_ precipitation. *A. porrum* leaf segments were transformed using a biolistic gun (Plant Industry CSIRO, Australia), at a helium pressure of approximately 400 kPa and under 550 mm of Hg vacuum and then incubated on MS media (Murashige and Skoog, 1962) in the dark for 22 hours.

### Confocal microscopy

GFP fusion proteins were visualised in hand sectioned soybean transgenic nodules or leaf segments of leek. Localisation of GFP fusion and tonoplast mCherry marker (vac-mCherry; Nelson *et al*., 2007) were visualised using a Pascal confocal laser scanning system (Carl Zeiss, Germany) attached to an Axiovert microscope (Carl Zeiss, Germany) or a Leica SP5 II confocal microscope (Australian Centre of Microscopy and Microanalysis, The University of Sydney). To visualise nodule membranes including the SM, sections were counterstained with FM4-64 (30 mg/ml; Life Technologies)(Limpens et al. 2009) for 1 hour on ice and mounted in 0.1 M PBS (pH 7.2). GFP (excitation, 488 nm argon laser; emission, 505/530), FM4-64 fluorescence (excitation, 561 nm diode pumped solid state laser; emission, 691/800), mCherry (excitation 543 nm laser emission 560 nm) were captured using Leica Microsystems LAS AF software and images processed using Fiji image analysis software (Schindelin *et al.*, 2012; Schneider *et al.*, 2012).

### Symbiosome and microsomal membrane isolation and proteomics

SM was isolated from nodules of 80 soybean plants at 26 and 27 DAI following previously documented protocols (Day *et al.*, 1989; Panter *et al*., 2000; Clarke *et al*., 2015). The SM fraction isolated from Percoll gradient-purified symbiosomes was resuspended in 1 M urea and used directly for proteomic analysis. This reduced the loss of protein during steps that remove lipid from the sample.

A microsomal membrane sample was prepared by carefully grinding nodules in ice-cold extraction buffer [25 mM MES-KOH pH 7.0, 350 mM mannitol, 3 mM MgSO_4_, 1 mM PMSF plus Complete, mini, EDTA-free protease inhibitor cocktail (Roche)]. The homogenate was filtered through 4 layers of Miracloth and centrifuged at 20,000 g for 20 min at 4°C, to pellet intact organelles and symbiosomes. The supernatant, likely to be enriched in plasma membrane, but also containing membrane fragments from ER and broken organelles, was centrifuged at 100,000 g for 1 hour at 4°C and the pellet was resuspended in 8M Urea. The pellet from the initial centrifugation, containing intact symbiosomes and other organelles, was resuspended in extraction buffer and vortexed to break the symbiosomes (Clarke et al., 2015). After centrifugation (20,000 g, 20 min, 4°C), the supernatant was collected and centrifuged as above to collect the SM-enriched pellet, which was resuspended in 8 M urea.

Protein concentration was determined using the Bradford Protein Assay (BioRad). Isolated membrane samples were analysed at the La Trobe Comprehensive Proteomics Platform (La Trobe University). Samples were digested with trypsin (Clarke etal. 2015) e cleaned using stage-tips preparations with 3 plugs of Empore polystyrenedivinylbenzene (SBD-XC) copolymer disks (Sigma Aldrich, MO, USA) for solid phase extraction following the manufacturers instructions.

For the analysis, peptides were reconstituted in 0.1% formic acid and 2% acetonitrile and loaded onto a trap column (C_18_ PepMap 100 μm i.d. × 2 cm trapping column, Thermo Fisher Scientific) at 5 µL/min for 6 min using a Thermo Scientific UltiMate 3000 RSLCnano system and washed before switching to the analytical column (BEH C_18_, 1.7 μm, 130 Å and 75 μm ID × 25 cm, Waters). The separation of peptides was performed at 45 °C, 250 nL/min using a linear ACN gradient of buffer A (0.1% formic acid, 2% ACN) and buffer B (0.1% formic acid, 80% ACN). Data were collected on a Q Exactive HF (Thermo-Fisher Scientific) in Data Dependent Acquisition mode using *m*/*z* 350–1500 as MS scan range at 60 000 resolution. HCD MS/MS spectra were collected for the 7 most intense ions per MS scan at 60 000 resolution with a normalized collision energy of 28% and an isolation window of 1.4 *m*/*z*. Dynamic exclusion parameters were set as follows: exclude isotope on, duration 30 s and peptide match preferred. Other instrument parameters for the Orbitrap were MS maximum injection time 30 ms with AGC target 3 × 10^6^, MSMS for a maximum injection time of 110 ms with AGT target of 1 × 10^5^.

Raw files consisting of high-resolution MS/MS spectra were processed with MaxQuant version 1.5.5.1 to detect features and identify proteins using the search engine Andromeda. Sequence data for soybean from Phytozome (https://phytozome.jgi.doe.gov/pz/portal.html#!info?alias=Org_Gmax) was used as the database for the search engine. To assess the false discovery rate (FDR) a decoy data set was generated by MaxQuant after reversing the sequence database. Theoretical spectra were generated using trypsin as the enzyme and allowing two missed cleavages. The minimum required peptide length used was seven amino acids. Carbamidomethylation of Cys was set as a fixed modification, while N-acetylation of proteins and oxidation of Met were set as variable modifications. Precursor mass tolerance was set to 5 ppm and MS/MS tolerance to 0.5 Da. The “match between runs” option was enabled in MaxQuant to transfer identifications made between runs on the basis of matching precursors with high mass accuracy. PSM and protein identifications were filtered using a target-decoy approach at a false discovery rate (FDR) of 1%.

### Yeast complementation assays

*Δccc1* yeast mutants in either a DY150 (*MATa ura3-52, leu2-3, 112, trp1-1, his3-11, 15, ade2-1, can1-100(oc) Δccc1*::HIS3) or BY4741 (*MATa, his3Δ1, leu2Δ0, met15Δ0, ura3Δ0*) background were transformed either by the lithium acetate method (described by Gietz et al. 1995) or via transformation of competent yeast cells (Dohmen *et al*., 1991).

Yeast containing the desired expression vectors were grown and OD600 adjusted to 1. Dilutions were spotted onto SD media containing either 2% [w/v] glucose or 2% [w/v] galactose and 0-20 mM FeSO_4_ to test for iron transport. Cultures were grown at 28°C for 5 days. Images were taken with the G:Box UV transilluminator (Syngene).

### Perls/DAB staining

Nodules were isolated 35 DAI from *L. japonicus* Gifu and *sen1-1* mutant plants inoculated with *M. loti*. The nodules were vacuum infiltrated with fixative solution (0.25% glutaraldehyde, 4% paraformaldehyde and 2.5% sucrose in 0.05 M potassium phosphate buffer pH 7.4) for 30 minutes and then incubated overnight at 4°C. Nodules were washed with potassium phosphate buffer pH 7.2, 2 times for 10 minutes. Nodule samples were dehydrated through an ethanol gradient and infiltrated and embedded in Technovit 7100 (Heraeus Kulzer), following the manufacturers instruction. Specimens were sectioned to 6-10 µm using a rotary microtome.

Perls/DAB staining was done according to the methods described by Roschzttardtz *et al*. (2013) on sections of whole nodules except all washes were repeated 3 times.

### Acetylene Reduction Assay

To assess nitrogenase activity, an acetylene reduction assay was performed on the whole root systems of 5 plants per transformation construct. Each root system was incubated in 10% acetylene at 28°C for 60 minutes. Gas samples were taken at intervals and compared to ethylene standards. A Shimadzu Gas Chromatograph 9AM (Shimadzu) was used to determine the relative concentration of acetylene and ethylene using 0.5 ml of gas. The peak area for both gases was recorded. The number of nodules per vial and their weight was recorded and the rate of acetylene reduction calculated.

### Bioinformatics and Phylogenetic analysis

*G. max, M. truncatula* and *A. thaliana* homologues to Nodulin21 (GmVTL1) and AtVIT1 were identified by selecting protein homologs on the gene page of Phytozome (http://phytozome.jgi.doe.gov/pz/portal.html#). BLASTP searches were used to identify VIT family members from *L. japonicus* in Lotus Base (Mun *et al*., 2016) except for LjSEN1 (BAL46698) which was obtained from NCBI. Sequences for characterised VIT family members were obtained based on published accession numbers from NCBI (http://www.ncbi.nlm.nih.gov/) or UniProt (http://www.uniprot.org/uniprot/). These included *S. cerevisiae* CCC1 (Sc, VIT1, P47818), *Centaurea cyanus* VIT1 (CcVIT1, W8VRG3), *Tulipa gesneriana* VIT1 (TgVIT1, BAH98154), rice OsVIT1 (Os04g0463400, BAS89575) and OsVIT2 (Os09g0396900, Q6ERE5.2)

A maximum-likelihood phylogenetic tree was produced for the VIT family using MEGA6.0 (Tamura *et al*., 2013). Amino acid sequences from characterised VIT family members [AtVIT1 (Kim et al. 2006), AtVTL1, AtVTL2, AtVTL5 (Gollhofer et al. 2014), OsVIT1, OsVIT2 (Zhang et al. 2012), TgVIT1 (Momonoi et al., 2012), CcVIT1 (Yoshida and Negishi, 2013) and LjSEN1 (Suganuma *et al*., 2003; Hakoyama *et al.*, 2012)] were aligned with all *G. max, M. truncatula, L. japonicus* and *A. thaliana* members using MUSCLE. The best substitution model was predicted to be the JTT model with gamma distribution. *S. cerevisiae* CCC1 was used as an out-group. The robustness of each node was assessed via 1000 bootstrap replicates.

## Results

### Phylogeny of the VIT family

Phylogenetic analysis of the plant VIT family identifies two separate groups (Fig. **1**). Group 1 is more closely related to *S. cerevisiae* CCC1 and includes the characterised ferrous iron transporters AtVIT1, CcVIT1, OsVIT1, OsVIT2 and TgVIT1. The second, larger group contains many uncharacterised members of the protein family, including GmVTL1 and GmVTL2, as well as the iron-responsive ferrous iron transporters of *A. thaliana*, AtVTL1, AtVTL2 and AtVTL5. AtVTL1 and AtVTL2 are closely related to each other and distant from AtVTL5. GmVTL1 and GmVTL2 are also closely related and are present in a legume specific clade that contains *L. japonicus* SEN1 and another nodule-expressed protein *M. truncatula* VTL8 (Fig. **1**).

**Figure 1.**
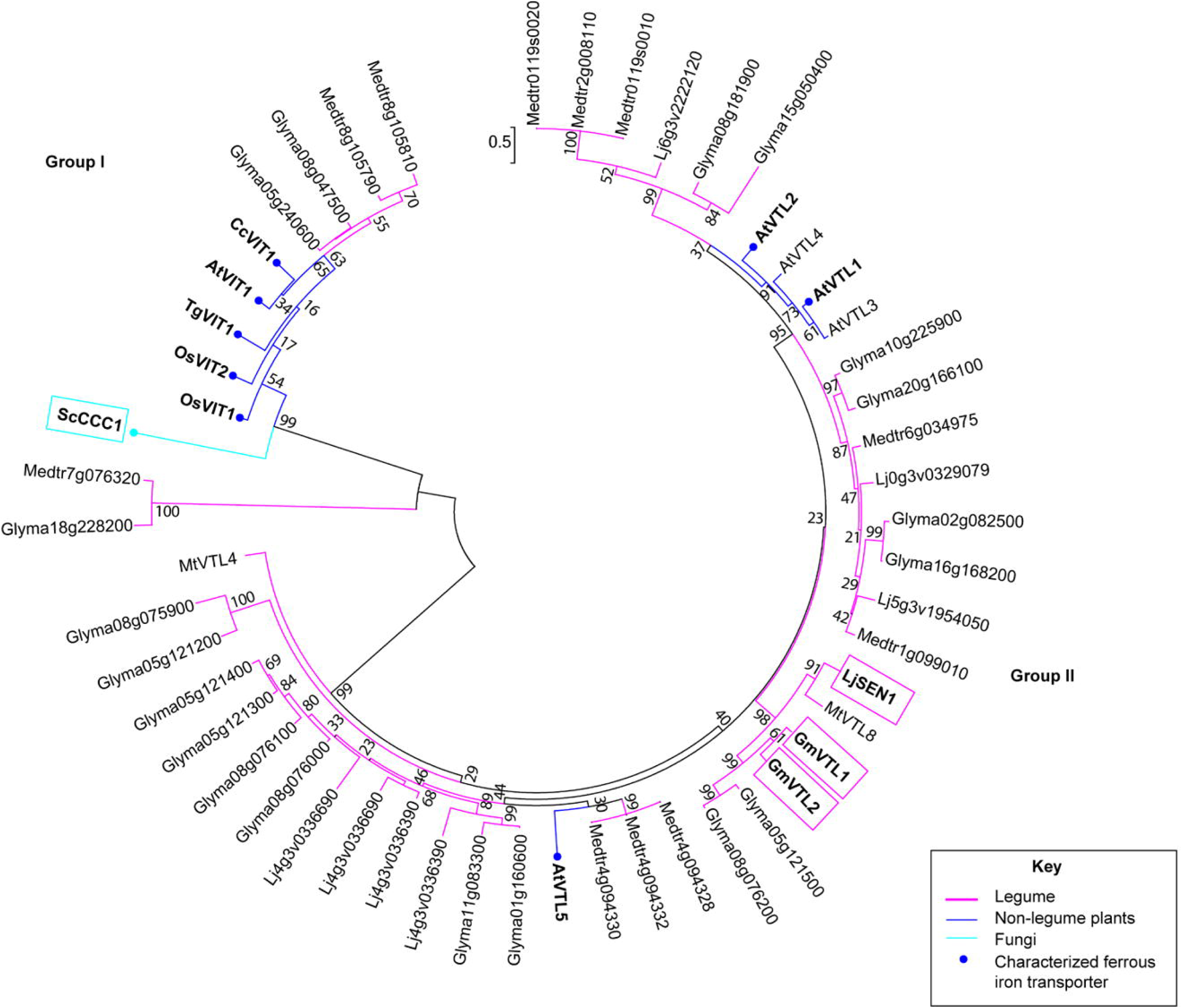
Phylogenetic analysis of characterised vacuolar iron transporter family (VIT) family members, together with all family members in *G. max, truncatula, L. japonicus* and *A. thaliana.* Sequences were aligned via MUSCLE in Mega6.0. The phylogenetic tree is a maximum likelihood tree using the JTT + G model, produced using Mega 6.0, with *S. cerevisiae* CCC1 (ScCCC1) as the out-group. The robustness of each node was assessed via 1000 bootstrap replicates. Nodes are labelled with their corresponding bootstrap value. Named sequences from *A. thaliana* are AtVIT1 (At2g01770), AtVTL1 (At1g21140), AtVTL2 (At1g76800), AtVTL3 (At3g43630), AtVTL4 (At3g43660), AtVTL5 (At3g25190), from *Centaurea cyanus* is CcVIT1 (BAO52026), from soybean are GmVTL1 (Glyma.05G121600, NP_001236825) and GmVTL2 (Glyma.08G076300, XP_003531056), from *L. japonicus* is LjSEN1 (BAL46698), from rice are OsVIT1 (Os04g0463400, BAS89575) and OsVIT2 (Os09g0396900, Q6ERE5.2), from *Tulipa gesneriana* is TgVIT1 (BAH59029) and from *M. truncatula* are MtVTL4 (Medtr4g094325) and MtVTL8 (Medtr4g094335). Proteins functionally characterised as ferrous iron transporters have a circle at their tip (Kim et al. 2006; Gollhofer *et al*., 2011; Gollhofer *et al*., 2014; Momonoi *et al*., 2009; Yoshida & Negishi, 2013; Zhang *et at*., 2012).

### GmVTL1 and GmVTL2 have enhanced expression in mature nodules, specifically in infected cells

Soybean transcriptome data shows *GmVTL1* and *GmVTL2* have significantly higher expression than any other VIT family members in nodules and more than 100-fold higher expression in nodules compared to roots (Severin *et al*., 2010; Libault *et al.*, 2010; Brear *et al.*, 2013). To confirm this and to assess temporal expression in the nodule, quantitative real time PCR analysis was performed. Expression of *GmVTL1* and *GmVTL2* was strongly enhanced in nodules compared to root tissue, with approximately 130 and 185 times higher expression in nodules than roots, respectively (Fig. **2a**). In nodules, 25 DAI expression of *GmVTL1* was 1.7 times higher than that of *GmVTL2*.

**Figure 2.**
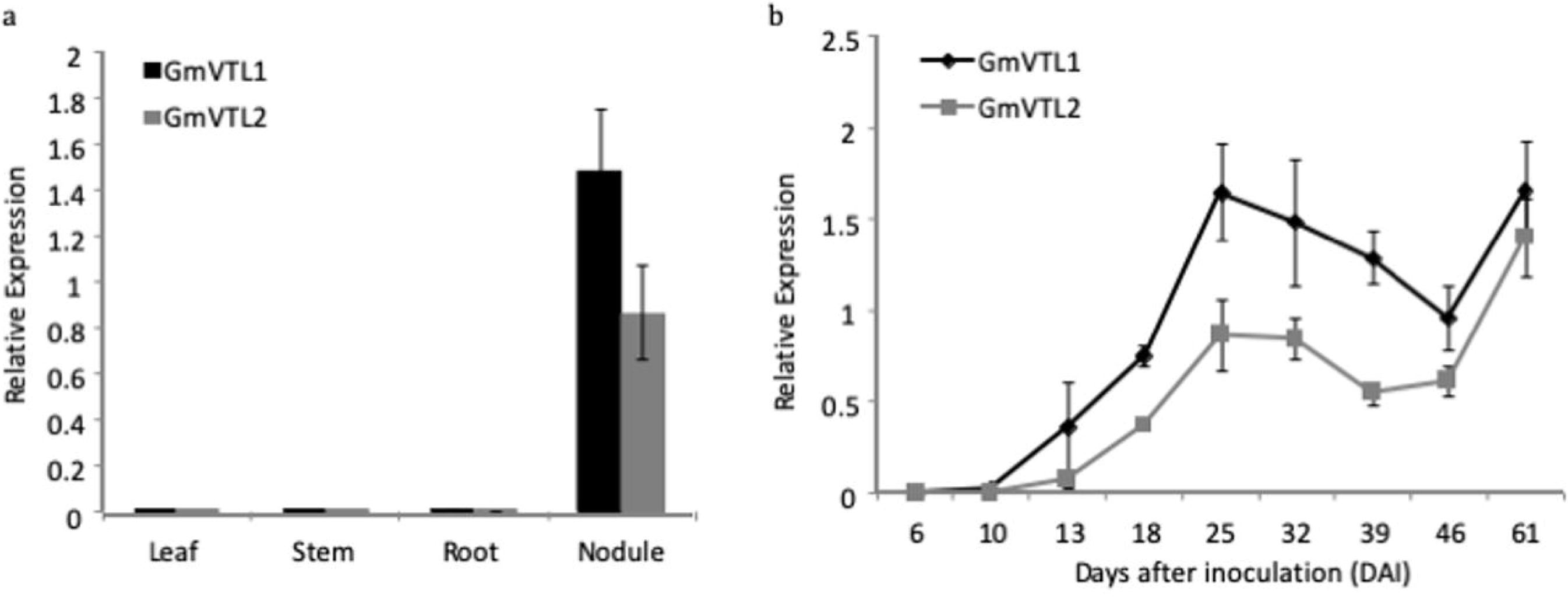
Relative expression of *GmVTL1* and *GmVTL2* in *G. max* tissues and throughout nodule development. **(a)** *GmVTL1* and *GmVTL2* expression in *G. max* leaf, stem, root and nodule tissue isolated 25 days after inoculation (DAI) with *B. japonicum.* **(b)** *GmVTL1* and *GmVTL2* expression during nodule development. At 6 & 10 DAI with *B. japonicum* the RNA used as template for cDNA synthesis was isolated from nodules and roots. For all other timepoints RNA was isolated from nodules. The time points when flowering and pod formation were occurring are indicated. The mean relative expression is shown for 6 biological replicates (except for 13 DAI, which is an average of 3 biological replicates) and error bars represent standard error. Expression was normalized to *G. max* ubiquitin, *GmUbi3* (Glyma.20G141600).

Expression of *GmVTL1* and *GmVTL2* was first detected at 13 DAI and continued to increase until 25 DAI (Fig. **2b**). This increase in expression coincided with increasing nitrogenase activity, measured by an acetylene reduction assay (Supporting Information Fig. S1). Expression subsequently declined and there was significantly lower expression by 46 DAI but this was followed by an increase at 61 DAI (Fig. **2b**). The highest level of expression was observed in 61 DAI nodules, which coincided with pod formation (Fig. **2b**). The expression pattern for both genes was similar during nodule development but expression of *GmVTL2* was always lower.

The localisation of *GmVTL1* expression in nodules was investigated using a promoter-GUS fusion (Fig. **3**). A 512 bp region upstream of the *GmVTL1* coding region activated the expression of *β-glucoronidase* in the large infected cells of the inner cortex and in cells surrounding the vasculature (Fig. **3**). A lower level of expression was present in the smaller uninfected cells, cells of the inner cortex and a layer of cells in the outer cortex (Fig. **3**). No GUS activity was observed in the root (results not shown).

**Figure 3.**
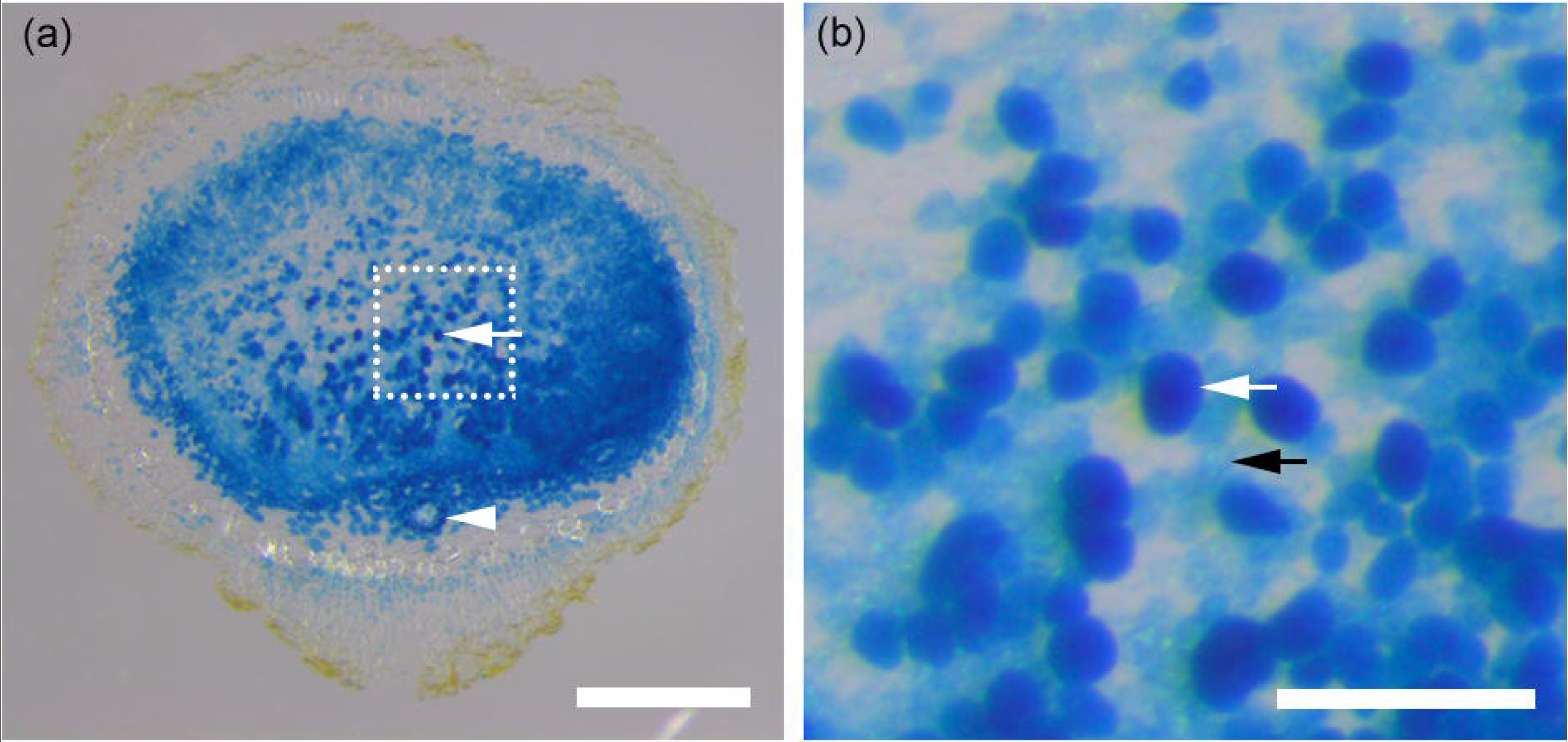
*GmVTL1* is expressed in infected cells and cells surrounding the vasculature in the *G. max* nodule. **(a)** A nodule expressing β-glucuronidase from the *GmVTL1* promoter. Infected cells have stained dark blue indicating activation of *GmVTL1* and subsequent expression of β-glucuronidase in these cells (white arrow). Cells surrounding the vasculature have also stained dark blue (arrowhead). **(b)** Magnification of the boxed regions presented in **(a)**. White arrows indicate infected cells stained dark blue, while black arrows indicate uninfected cells with pale blue staining. The scale bars represent 500 μm in **(a)** and 125 μm in **(b)**

### GmVTL1 and GmVTL2 localise to the vacuole in leek epidermal cells

The localisation of GFP fusion constructs for both GmVTL1 and GmVTL2 was tested in leek epidermal cells using particle bombardment. GmVTL1-GFP, GFP-GmVTL1 and GmVTL2-GFP colocalised with the tonoplast marker mCherry-vac (Nelson *et al*., 2007) (Fig. **4**).

**Figure 4.**
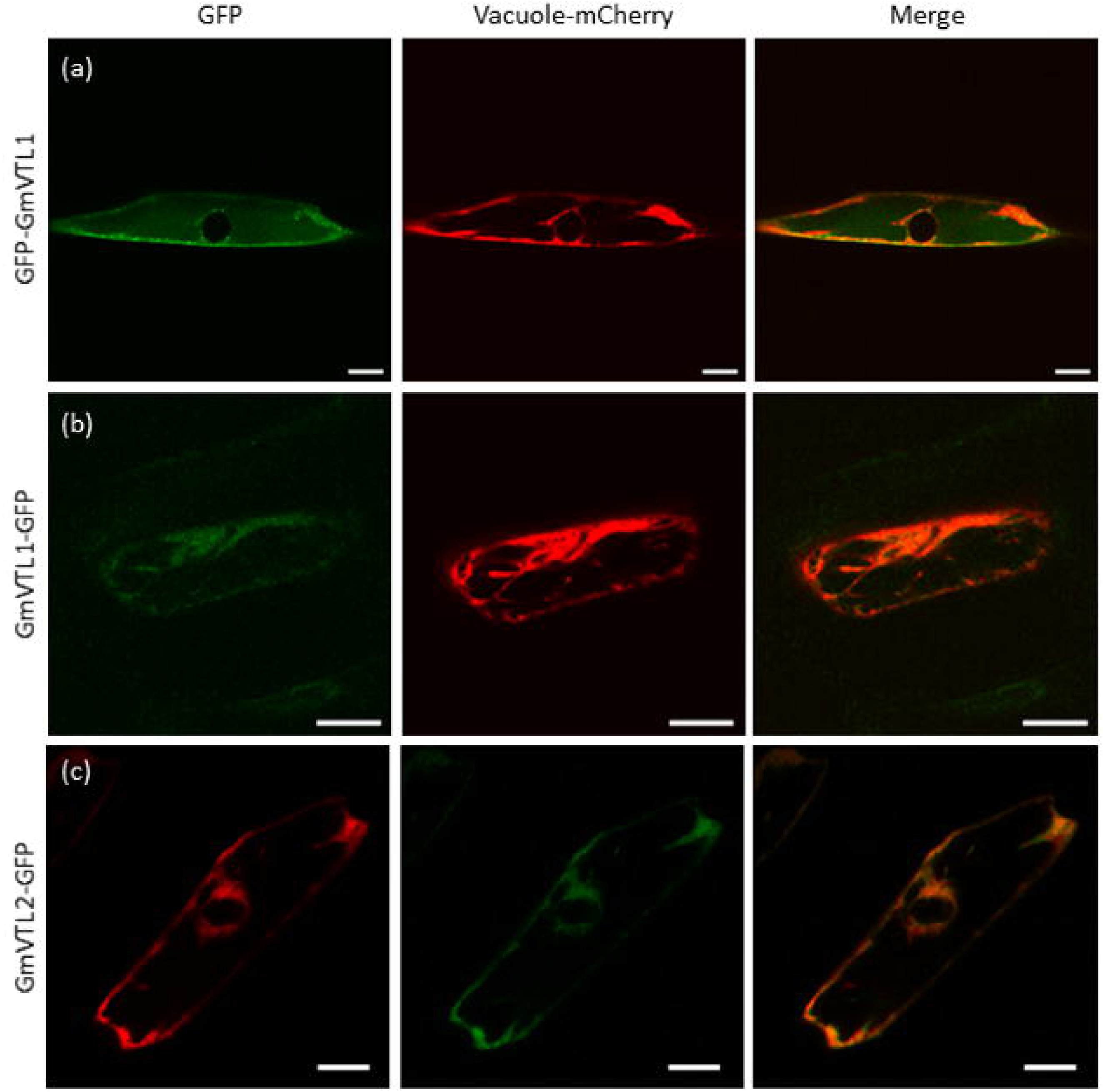
Co-localisation of GFP-fusions of GmVTL1 and GmVTL2 signal with Vac-mCherry in leek epidermal cells. Epidermal cells co-expressing (a) GFP-GmVIT1, (b) GmVIT1-GFP or (c) GmVIT2-GFP (images to the left) and the vacuolar membrane marker, Vac-mCherry (central images). Overlays of the red and green channels are shown to the right. All images were taken at a single focal plane. The scale bars represent 20 μm.

### GmVTL1 localises to the SM

The subcellular localisation of GmVTL1 in infected cells was investigated by fusing the green fluorescent protein (GFP) marker to the N-terminus of GmVTL1 and expressing this transgene fusion in nodules produced on transformed hairy roots. In infected cells of nodules 22 DAI, GmVTL1 localised to ring like structures that overlapped with the FM4-64 stained SM (Fig. **5**). Analysis of the intensity of the FM4-64 and GFP signals confirmed that the GFP and FM4-64 signals overlapped (Fig. **5**).

**Figure 5.**
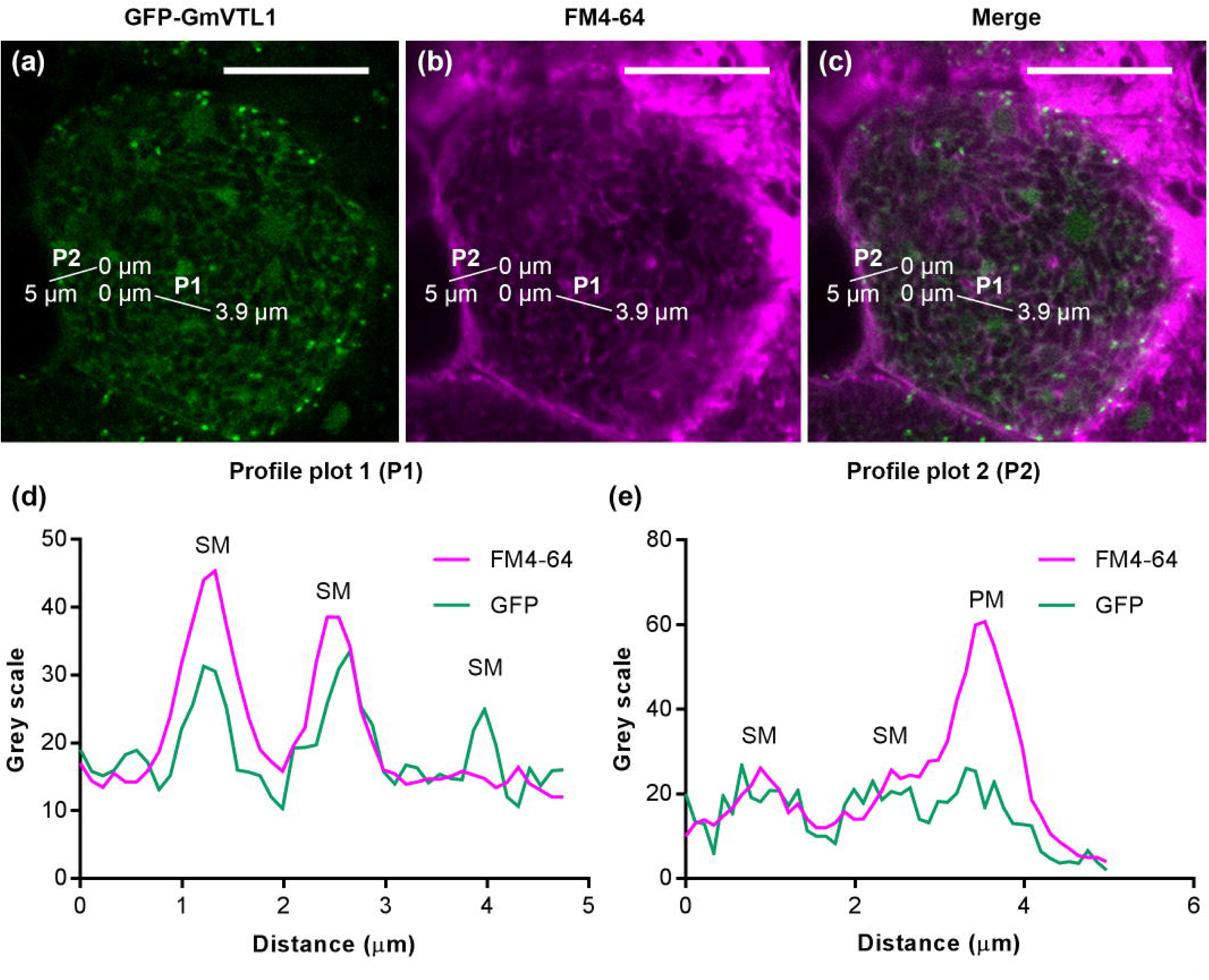
Co-localisation of GFP-GmVTL1 signal and FM4-64 membrane signal in infected cells of *G. max* nodules. GmVTL1 localisation in an infected cell **(a)**, membrane stained with FM4-64 **(b)** and overlay of the magenta (FM4-64) and green channels (GmVTL1-GFP)**(c)**. The co-localisation of the two signals was assessed by plotting the grey scale profile across two lines, P1 and P2, starting at 0 μm. Co-localisation can be seen in the resulting profile plots 1 **(d)** and 2 **(e)**, when peaks overlap. The overlapping peaks are at the symbiosome membrane (SM). The scale bars represent 12 μm.

Localisation of GmVTL2 in infected nodules cells using GFP fusions was not as definite as it was for GmVTL1. GFP aggregates could be seen overlapping FM4-64 in some parts of cells on what could be SM but the results were not as clear (results not shown).

The SM localisation of GmVTL1 was confirmed by identification of peptides corresponding to the protein in purified SM samples using proteomics (Table **1**). GmVTL2 was also identified on the SM in this analysis. Five unique peptides for GmVTL1 and three unique peptides for GmVTL2 covered 25% of the amino acid sequence of each protein in one SM sample (Table **1**). The peptides were also detected in a second sample, but were less abundant (Table **1**). A peptide matching GmVTL2 was also identified in the previously published SM proteome (Clarke *et al*., 2015).

**Table 1.**
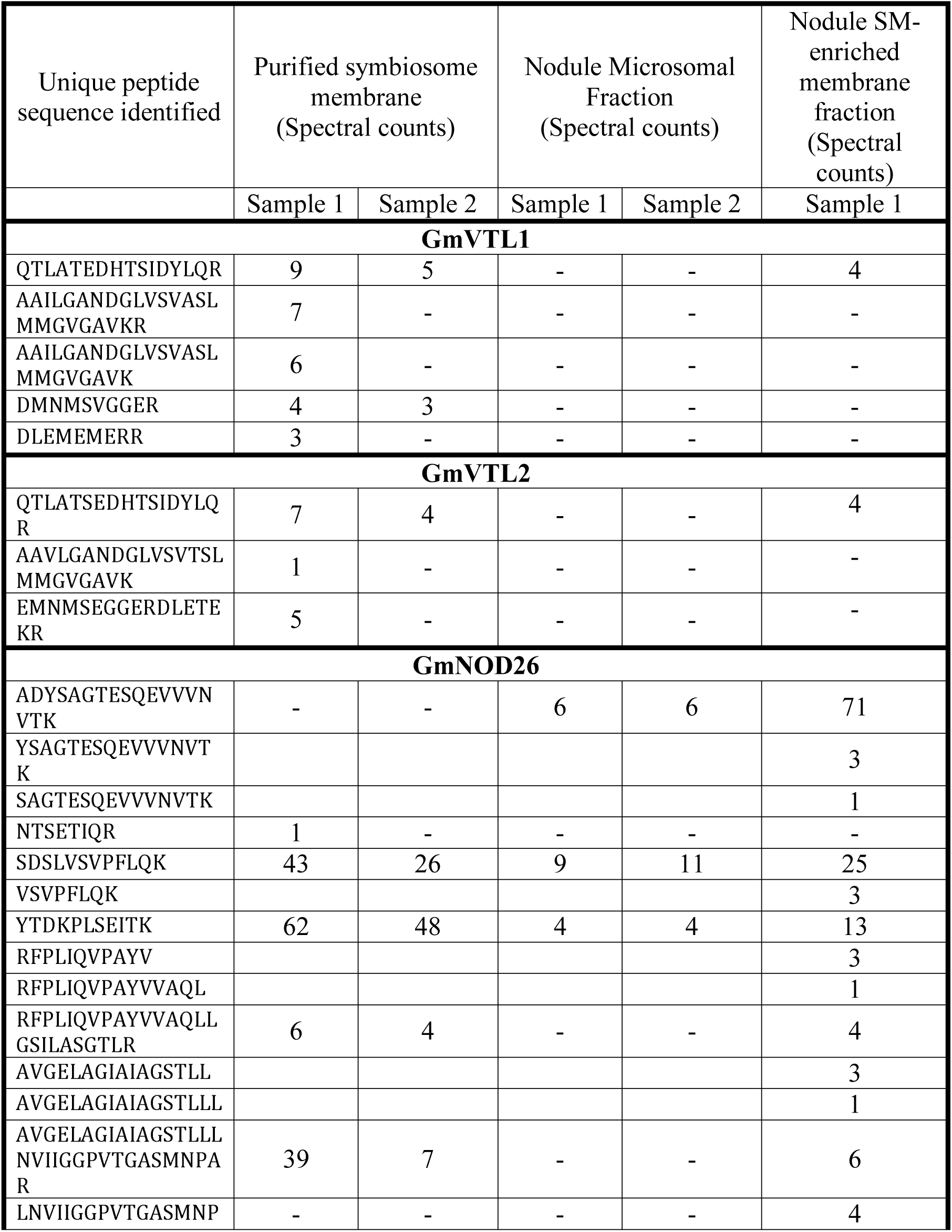
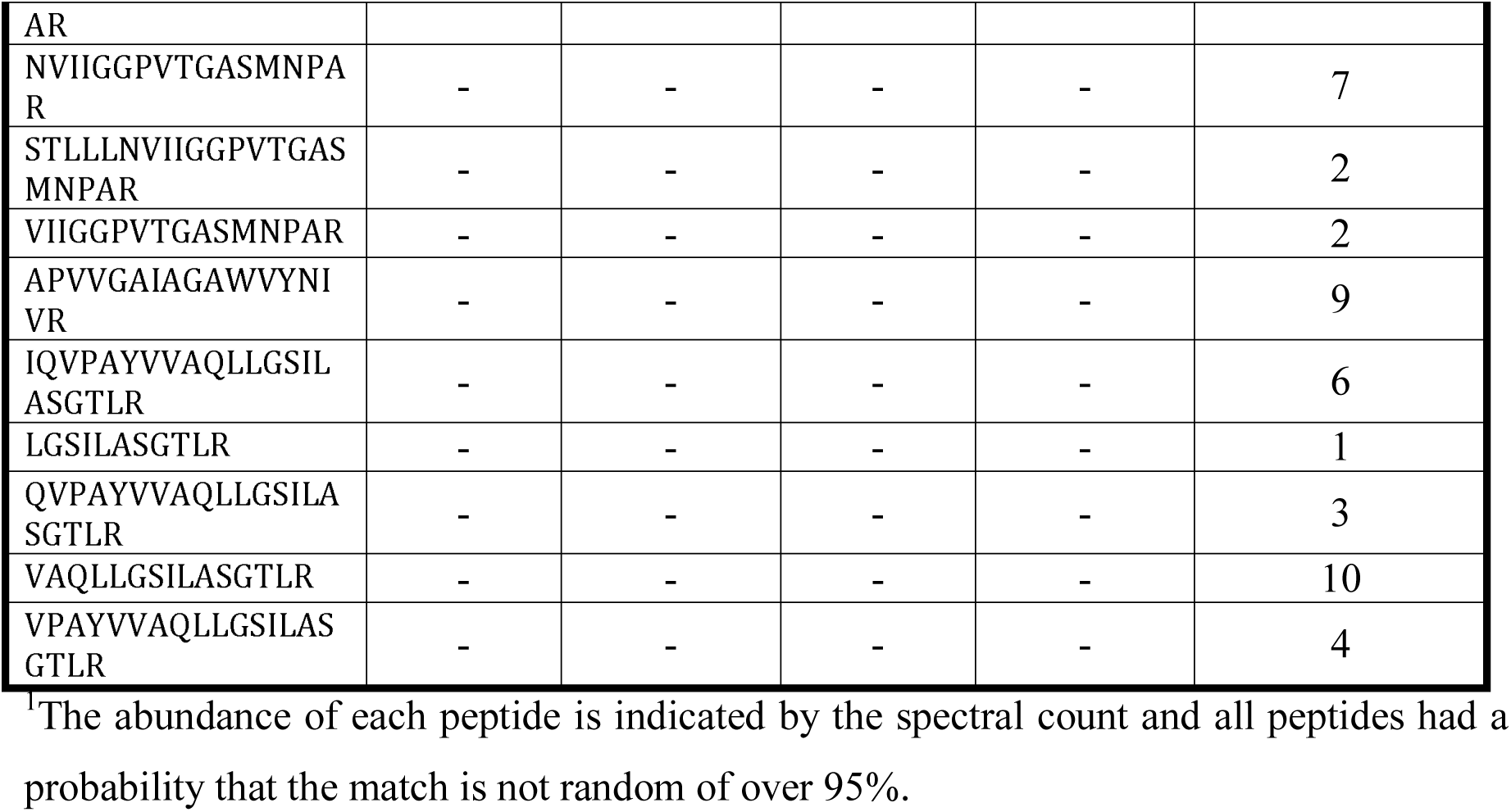
Unique GmVTL1, GmVTL2 and GmNOD26 peptides^1^ identified in purified symbiosome membrane, a microsomal membrane fraction, and a symbiosome-enriched membrane sample from soybean nodule homogenate. See methods for details of membrane preparation.

To show that there was enrichment of GmVTL peptides in the SM, we prepared a microsomal fraction from material with symbiosomes removed (termed plasma membrane (PM)-enriched) as well as an SM-enriched fraction from a soybean nodule homogenate, and compared them to purified SM, using the abundance of peptides corresponding to GmNOD26, a well characterised SM-specific protein (Wallace et al. 2006; Clarke et al. 2015), as an SM marker (Table **1**). NOD26 peptides were approximately six times more abundant in the SM sample compared to the microsomal membrane sample. In addition, only small numbers of spectra of the peptides were present in the microsomal fraction, while NOD26 identified in the purified SM and SM-enriched sample was represented by a large number of spectra. Peptides corresponding to both GmVTL1 and 2 were identified in the purified SM and SM-enriched sample but were not detected in the microsomal sample (Table **1**). This strongly suggests that the two VTL proteins are located on the SM.

### GmVTL1 and 2 complement the yeast ferrous iron transport mutant *Δccc1*

Transport into the symbiosome is directionally similar to transport out of the cell or into an organelle, the same direction in which CCC1 transports ferrous iron to sequester it in the yeast vacuole (Fu *et al*., 1994; Lapinskas *et al*., 1996; Li *et al*., 2001). The *Δccc1* yeast mutant is unable to survive on media containing high concentrations of iron and has been used in complementation experiments to confirm iron transport by a number of plant VIT family members (Kim *et al*., 2006; Momonoi *et al*., 2009; Zhang *et al*., 2012; Gollhofer *et al*., 2014; Connorton et al., 2017).

*GmVTL1, GmVTL2* and *Lotus japonicus SEN1* (*LjSEN1*, an orthologue of the proteins encoded by these genes) were cloned into the yeast expression vector pDR196-Gateway and transformed into two *Δccc1* yeast strains. On media with a low iron concentration both *Δccc1* yeast containing the empty vector (EV-*Δccc1*) and wild-type cells grew to the same degree, but on media containing 10 mM FeSO_4_ the EV-*Δ ccc1* yeast could not survive (Fig. **6**). Transformation of *Δccc1* DY150 with *GmVTL1* rescued its growth on high iron, suggesting that GmVTL1, like AtVIT1 (Kim et al. 2006), can transport iron into yeast vacuoles. G*mVTL2* also was able to rescue the growth of the *Δccc1* DY150 yeast on media containing 2, and to some extent, 5 mM iron (Fig. **6**). However, LjSEN1, like the empty vector, was not able to do this (Fig. **6**). Similar results were obtained in the *Δccc1* BY4741 yeast, except that *GmVTL2* was more effective in this strain (Supporting Information Fig. S2). Our results suggest that GmVTL2 has some ability to transport iron, although it is not as effective as GmVTL1.

**Figure 6.**
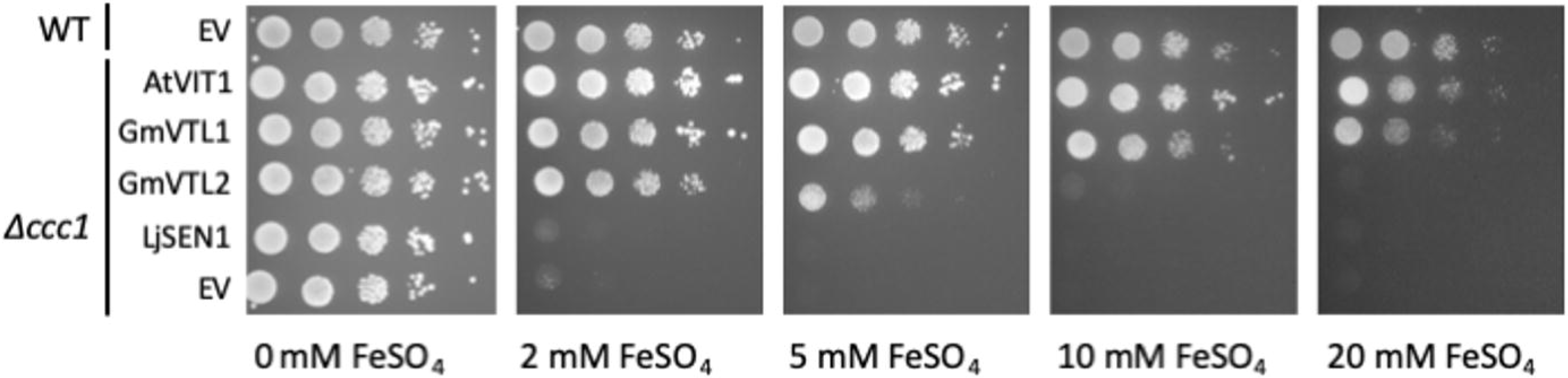
GmVTL1 and GmVTL2 transport iron into the vacuole in the DY150 yeast iron transport mutant *Δccc1*. Wild type (WT) DY150 yeast was transformed with the empty vector plasmid (EV; pDR196-GW. The Δccc1 yeast strain derived from DY150 was transformed with pDR196-GW vector containing the full-length coding sequence of *A. thaliana* VIT1 (AtVIT1, positive control), *G. max* VTL1 (GmVTL1), *G. max* VTL2 (GmVTL2), and EV, under control of the constitutive PMA promoter. Serial dilutions of each yeast transformant were applied to SD plates (that include 1.6 µM FeCl_3_) with the addition of 0, 2, 5, 10 or 20 mM FeSO_4_ and the plates grown for 3 days.

### *L. japonicus sen1* is unable to complement the *Δccc1* yeast mutant due to mislocalisation

The inability of LjSEN1 to complement *Δccc1* yeast suggests either that it doesn’t transport iron or that it is misdirected or inactive in the yeast system. We used GFP-fusions to test the localisation of the LjSEN1 protein and compared it with that of GmVTL1 when expressed in yeast. The tonoplast of the yeast cells was stained with FM4-64 to better visualise co-localisation. The results clearly show that GmVTL1 co-localised with the FM4-64 stain on the tonoplast (Supporting Information Fig. S3). LjSEN1, on the other hand, appeared to localise only within the vacuolar lumen, rather than the tonoplast membrane (no obvious co-localisation with the FM4-64 stained tonoplast; Supporting Information Fig. S3). This mis-localisation of LjSEN1 is the most likely explanation for its inability to complement the yeast mutant but reduced expression due to inclusion of codons that are not optimal for expression in yeast is another explanation.

One obvious difference between LjSEN1 and GmVTL1 is that the former is missing an N-terminal extension evident in GmVTL1 (Supporting Information Fig. S4). We investigated whether addition of the first seven amino acids of GmVTL1 to LjSEN1 would improve iron transport in yeast. However, when the construct expressing this fusion was expressed in the *Δccc1* mutant, the yeast grew no better than those expressing wild type SEN1 (results not shown) suggesting that the extension makes no difference to the localisation of the SEN1 protein in yeast.

### Mutations that block nitrogen fixation in *Ljsen1* mutants also block iron transport in yeast

Since *LjSEN1* could not complement the *Δccc1* mutant, we investigated a possible link between the mutations that block nitrogen fixation in *sen1* nodules and its ability to transport iron. There were a number of point mutations characterised in *sen1* that caused a reduction or complete blockage of N-fixation (Hakoyama et al, 2012). In *sen1-1* a C122T nucleotide substitution results in an A41V amino acid substitution; in *sen1-2* a G332A substitution results in an R111K amino acid substitution; and in *sen1-5* a G572A produces a G191E substitution (an alignment showing the amino acids which were substituted is shown in Fig. S4). All the mutated nucleotides are conserved in GmVTL1 and GmVTL2. To investigate whether the same mutations in GmVTL1 affect its iron transport capacity, we generated mutated versions of GmVTL1 gene to create GmVTL1-A54V, -R124K and - G202K, and expressed them in the *Δccc1* yeast mutant (Supporting Information Fig. S4). The A54V and G202K substitutions prevented GmVTL1 from rescuing growth on media containing 2 mM (results not shown) and 5 mM FeSO_4_ (Fig. **7**). GmVTL-R124K grew almost as well as the unmutated GmVTL1, but at 10 mM FeSO_4_ there was minimal growth of the *Δccc1* expressing this mutated GmVTL1 (Fig **7**). Clearly, all of the mutations reduce the efficiency of iron transport by the protein.

**Figure 7.**
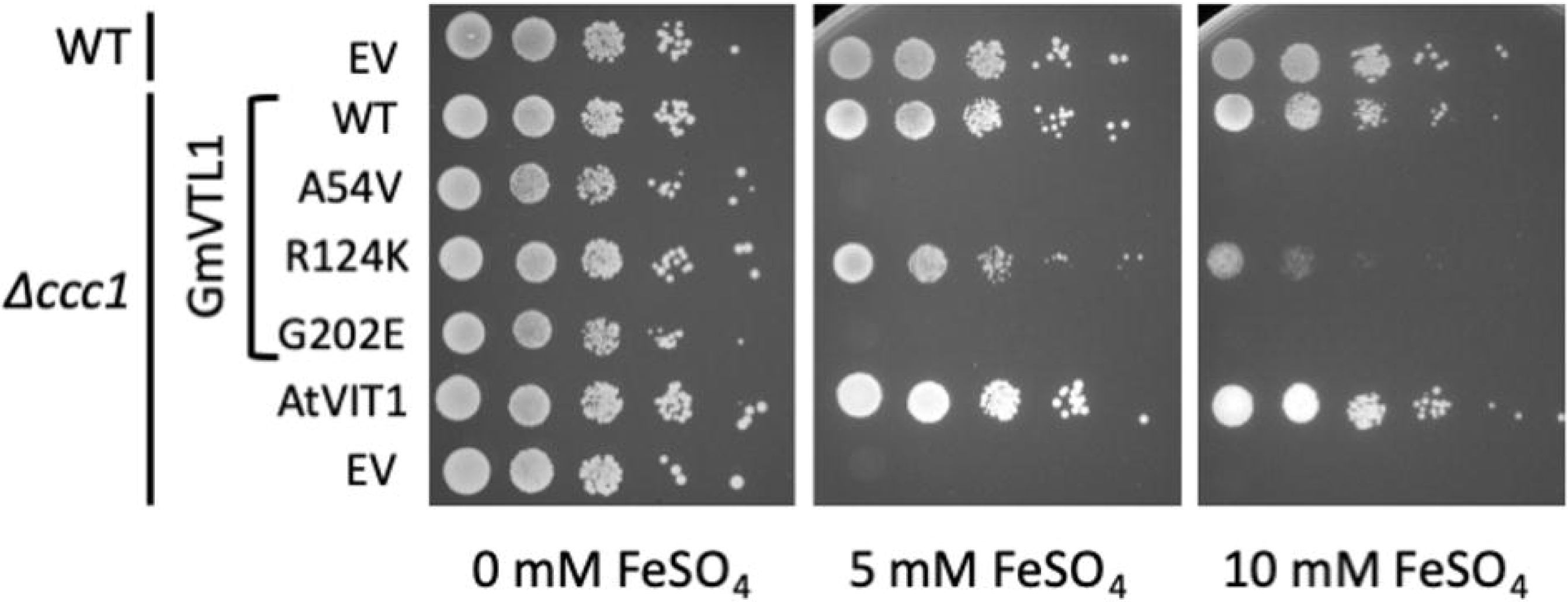
Mutations in GmVTL1 that block N-fixation in *Ljsen1* block iron transport into the vacuole in DY150 yeast mutant *Δccc1*. Wild type (WT) DY150 yeast was transformed with the empty vector plasmid (EV; pDR196-GW. The *Δccc1* yeast strain derived from DY150 was transformed with pDR196-GW vector containing the full-length coding sequence of *G. max VTL1* (GmVTL1), and three constructs incorporating the same mutations in *L. japonicus sen1-1* (A54V in GmVTL1), *sen1-2* (R124K) and *sen1-5* (G202E), *A. thaliana VIT1* (AtVIT1, positive control), and EV, under control of the constitutive PMA promoter. Serial dilutions of each yeast transformant were applied to SD plates (that include 1.6 µM FeCl_3_) with the addition of 0 mM or 10 mM FeSO_4_ and the plates grown for 3 days.

These results suggest that LjSEN1 does indeed transport iron *in planta*, but mislocalisation in yeast prevents it from complementing the *Δccc1* strain. To link the yeast transport results with the effects of the mutation in *Ljsen1-1* we stained nodules from the mutant and wild-type Gifu plants with Perls/DAB to assess iron localisation. Infected cells in wild-type nodules showed much stronger staining than in *sen1-1* infected cells suggesting that SEN1 plays a role in iron transport in infected cells (Supporting Information Fig. S5).

### GmVTL1 complements the *L. japonicus sen1-1* mutant

Our attempts to reduce expression of *GmVTL1* using hairy root transformation with RNAi constructs in soybean were unsuccessful with no reduction in expression (data not shown). We therefore used transformation of the *L. japonicus sen1* mutant to further investigate the role of GmVTL1.

The *Ljsen1-1* ethyl methanesulfonate (EMS) mutant has retarded growth under symbiotic conditions and is characterised by the presence of numerous small, white nodules that are unable to fix nitrogen throughout development (Suganuma *et al*., 2003). Nodules in these plants develop normally and their infected cells contain bacteria encapsulated by a SM, but they appear to senesce prematurely (Suganuma *et al.*, 2003). Like GmVTL1, LjSEN1 is expressed specifically in nodule infected cells, but its localisation in these cells was not determined (Suganuma *et al*., 2003; Hakoyama et al., 2012).

To investigate the *in planta* function of GmVTL1 we asked whether it could replace the function of LjSEN1 in *Lotus japonicus* nodules. Hairy root transformation was used to introduce an N-terminal-GFP fusion with the GmVTL1 coding sequence into the *sen1-1* mutant, under the control of the soybean leghemoglobin promoter (Gmlbc3) to drive expression in infected cells.

In *L. japonicus sen1-1* plants with transgenic nodules expressing GmVTL1 (*sen1*-*VTL1*), leaves were green and healthy in contrast to the *sen1-1* mutant transformed with the empty vector (*sen1*-EV), which had yellow leaves, a sign of nitrogen deficiency (Fig. **8a**, images for 5 complemented plants are shown in Supporting Information Fig. S6). Nodules from *sen1*-*VTL1* plants were pink, similar to wild-type nodules, highlighting the presence of leghemoglobin, in contrast to the white nodules of *sen1*-EV plants (Fig. **8a** and Supporting Information Fig. S6). While on average the *sen1*-*GmVTL1* plants had fewer nodules than *sen1*-EV or wild-type plants transformed with empty vector, they were substantially bigger (double the weight: Fig. **8**).

**Figure 8.**
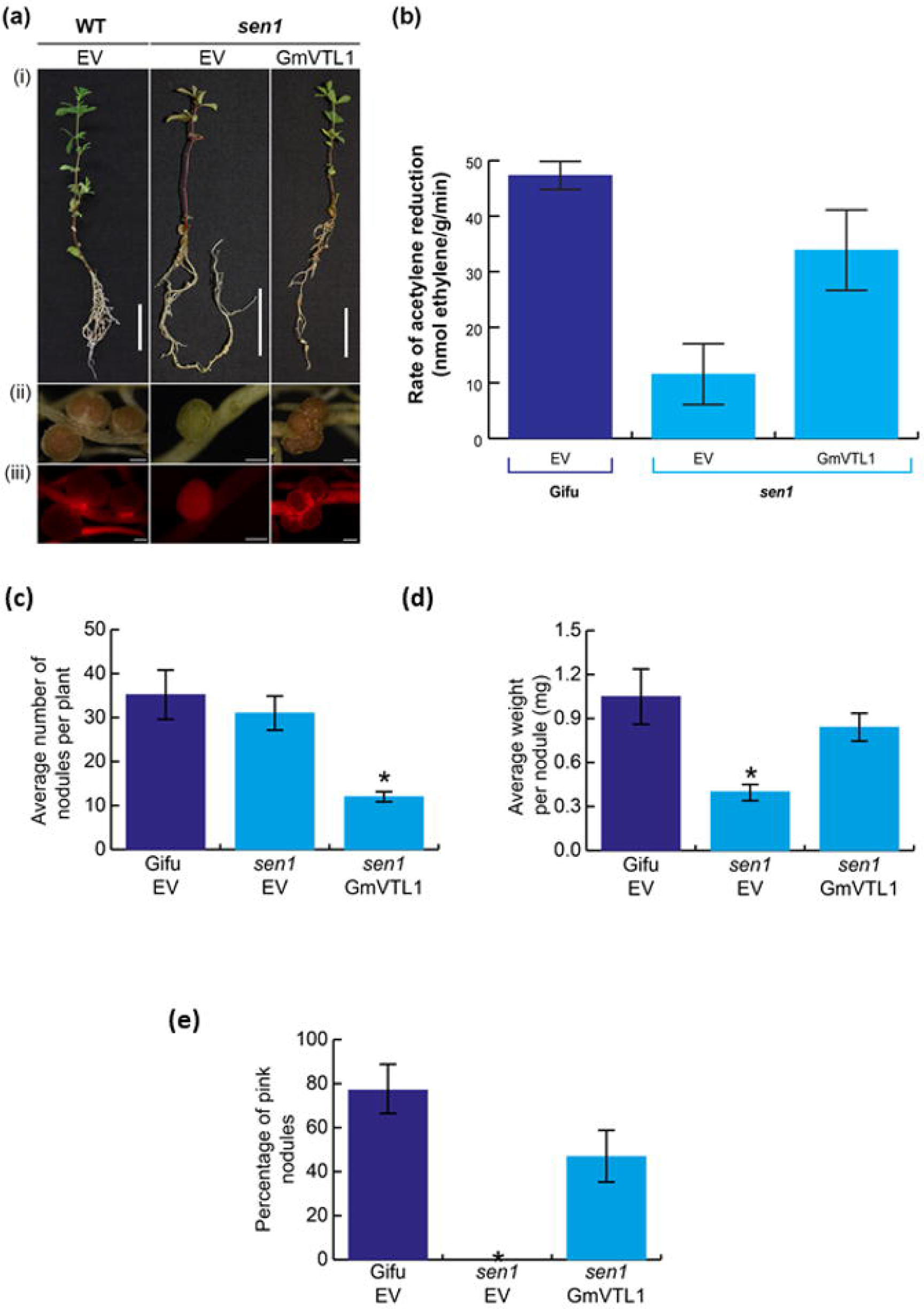
Expression of the *G. max VTL1* coding sequence in the *L. japonicus sen1-1* mutant complements the mutation and restores nitrogen fixation. **(a)** Phenotype of wild type Gifu plants transformed with empty vector (EV) (images to left), *sen1-1* mutant expressing empty vector (EV) (central images) and *sen1-1* mutant expressing *GmVTL1* coding sequence (*GmVTL1*) (images to right). Nodules present on transgenic hairy roots were identified 30 DAI with *M. loti* by identifying roots expressing DS red fluorescent protein. Phenotype of **(i)** whole plant and **(ii)** nodules. **(iii)** Image of each nodule showing expression of DSred fluorescent protein. The scale bars represent 2 cm **(i)** and 500 µm **(ii, iii). (b)** Average rate of acetylene reduction (nmol ethylene/g/min) for wild type *L. japonicus* Gifu transformed with the empty vector (Gifu-EV), *L. japonicus sen1-1* transformed with EV (LjSen1-EV) and *L. japonicus sen1-1* transformed with *GmVTL1* coding sequence (LjSen1-VTL1) expressed from the *G. max* leghemoglobin promoter. Averages were calculated from 5 plants per construct. Standard error bars are presented. Statistically significant differences were observed in a one-way ANOVA comparing the average rate of acetylene reduction (nmol ethylene/g/min) between sen1-EV and wild type Gifu-EV (p=0.002), as well as sen1-EV and sen1-GmVTL1 (p=0.03). **(c)** Average number of nodules per plant, Gifu-EV (n=5), *sen1*-EV (n=5) and *sen1*-GmVTL1(n=5). Standard error bars are presented. A P value < 0.002 (Student’s *t*-test) was obtained when comparing the average number of nodules per *Ljsen1-1* mutant transgenic plant transformed with either the empty vector (control) or the *GmVTL1* coding sequence. When comparing the *Ljsen1-1* mutant transformed with the GmVTL1 coding sequence and Gifu transformed with the empty vector a P value < 0.004 (Student’s *t*-test) was obtained. **(d)** Average weight per nodule on Gifu-EV (n=5), *sen1*-EV (n=5) and *sen1*-*GmVTL1* (n=5). Standard error bars are presented. A P value < 0.005 (Student’s *t*-test) was obtained when comparing *L. japonicus sen1-1* mutants transformed with either empty vector or *GmVTL1* coding sequence. **(e)** Average percentage of pink nodules per plant, Gifu-EV (n=5), *sen1*-EV (n=5) and *sen1*-*GmVTL1* (n=5). Standard error bars are presented. A P value < 0.005 (Student’s t-test) was obtained when comparing *L. japonicus sen1-1* mutants transformed with either empty vector or *GmVTL1* coding sequence

Nitrogenase activity (assessed by acetylene reduction) in *sen1*-EV nodules was much lower than in the empty vector transformed wild-type (Gifu-EV), indicating that the symbiosis was severely compromised (Fig. **8b**). Transformation of *sen1-1* plants with *GmVTL1* restored nitrogenase activity to wild-type levels (Fig. **8b**).

## Discussion

GmVTL1 and 2 are members of the VIT family but are only distantly related to the well-characterised vacuolar iron transporters, AtVIT1 and ScCCC1, grouping more closely with the iron-responsive AtVTL transporters (Fig. **1**). GmVTL1 and 2 are members of a larger soybean gene family (Brear *et al*., 2013, Cao, 2019), but are the only members that have greatly enhanced expression in nodules, consistent with a role in the symbiosis. Transcriptome data indicates that they may also be expressed in root hairs (Phytozome v10.3 expression data produced by Gary Stacey, University of Missouri; Cao, 2019). Their sequence similarity to LjSEN1, which is essential for the symbiosis (Suganuma *et al*., 2003; Hakoyama *et al*., 2012), suggests that they are the SEN1 orthologues in *G. max*.

Appearance of transcripts of *GmVTL1* and *GmVTL2* coincided with the onset of nitrogen fixation and increased in concert with the rate of nitrogen fixation during nodule maturation, suggesting that the two proteins are required in the mature, nitrogen-fixing nodule. A fusion between the GmVTL1 promoter and GUS suggested that expression of GmVTL1 is largely confined to the infected cells of the nodule. The pattern of expression for *GmVTL1* and *GmVTL2* is very similar to that reported for *LjSEN1* (Hakoyama *et al*., 2012) and is also similar to that for *L. japonicus Sst1* and *G. max ZIP1*, which encode transporters that mediate sulphur and zinc transport into the symbiosome, respectively (Wienkoop & Saalbach, 2003; Krusell *et al*., 2005; Moreau *et al*., 2002). Since both iron and sulphur are required by the bacteroids for nitrogen fixation, it is expected that their respective transporters are co-expressed (Krusell *et al*., 2005). After a decrease between 25 and 46 days after inoculation (DAI), both GmVTL1 and GmVTL2 showed a further increase in expression at 61 DAI, suggesting that they may also play a role during nodule senescence.

We used a combination of localisation of GFP fusions in leek cells and soybean nodules, together with proteomic analyses, to investigate the sub-cellular localisation of GmVTL1 and 2. Since symbiosomes largely replace vacuoles in soybean nodule mature infected cells (Bergersen 1982; Gavrin et al. 2014), in cells that do not contain symbiosomes, localisation to vacuoles is a good proxy for symbiosome localisation in nodules. Indeed, in Medicago some proteins that are normally localised on the vacuole are retargeted to the symbiosome (Gavrin et al. 2014) indicating that the mechanisms for targeting are likely to be shared. Both GmVTL1 and GmVTL2 were targeted to the tonoplast in leek epidermal cells (Fig. **3**), and also in yeast since they complemented the *Δccc1* mutant. The results for localisation of GFP-GmVTL1 (Fig. **4**) fusions in nodule infected cells were more robust than for GmVTL2 (results not shown), but peptides for both were identified in purified SM and symbiosome-enriched membrane samples, but not in a plasma membrane-enriched microsomal fraction (Table **1**). Together, these results give strong support to our proposal that GmVTL1 and 2 are localised on the SM in soybean.

Both GmVTL1 and GmVTL2 were able to complement the *Δccc1* yeast mutant confirming that they are able to transport iron. Although LjSEN1 could not rescue the yeast mutant, this was because the protein did not appear to localise to the yeast tonoplast in yeast. This makes definitively assigning an iron transport function to LjSEN1 difficult, but the DAB-staining of *Ljsen1* nodules, together with the ability of GmVTL1 to restore nitrogen-fixing capability in the mutant, makes it highly likely. Adding to this, we showed that GmVTL1 mutations mimicking the amino acid substitutions in *SEN1* that cause a dysfunctional symbiosis (Hakoyama et al., 2012), \ reduce iron transport when expressed in yeast. These results link the mutations that abolish nitrogen fixation in SEN1 to the function of GmVTL1 as an iron transporter.

The ability of *GmVTL1* to complement the *sen1* mutant, allowing formation of functional nodules, indicates that GmVTL1 and LjSEN1 play a similar role and suggests that GmVTL1 is important for proper functioning of the symbiosis. Other VIT family members, EgVIT1 and PfVIT, operate via a H^+^-antiport mechanism (Labarbuta et al., 2017, Kato et al. 2019). This mechanism is ideal for the transport of iron into symbiosomes, as the SM is energised by the action of P-type ATPases resulting in an acidic interior and a membrane potential positive on the inside of the symbiosome (Udvardi and Day, 1997). GmDMT1, another iron transporter localized to the SM (Kaiser et al. 2003), is an NRAMP family member and therefore likely to be a H^+^-symporter (González-Guerrero et al. 2016), making it an unlikely candidate for import of iron into the symbiosome. It is more likely to export stored iron from symbiosomes.

Although they both transport iron in yeast, GmVTL1 and 2 lack part of the cytoplasmic metal binding domain, including the conserved M149, important for ion binding in EgVIT1 (Kato et al. 2019). This suggests a unique mechanism for metal ion binding in the VTLs. Our work has identified three amino acids important for transport of iron by GmVTL1 and likely LjSEN1. Two, A54V and G202K, completely abolish complementation of the *Δccc1* mutant and a third, R124K, significantly reduces the efficiency of complementation compared to the wild type protein. R124 is conserved in most characterised VIT and VTL proteins and is adjacent to a conserved amino acid (E102 in EgVIT1; E in GmVTL2 and LjSEN1; D in GmVTL1) involved in ion binding in the metal binding domain (MBD) of EgVIT1 (Kato et al. 2019), suggesting that substitution in GmVTL1 may reduce efficiency of iron binding during transport. A54 and G202 are conserved in most VTL proteins but not in AtVIT1, EgVIT1 or PfVIT1. In EgVIT1 E32 is important for ion translocation from the MBD to the transmembrane domain. If the equivalent amino acid in GmVTL1 (Q55) has a similar role, this might explain why A54 has such a significant effect on complementation of the *Δccc1* mutant in yeast. These amino acids are candidates for further study to determine how the structure and transport of metals by VTL proteins differs from that of VIT proteins.

Together, these results support our proposal that GmVTL1 and SEN1 are able to transport iron *in planta* and play an important role in providing iron to the symbiont. However, while it is not unusual to observe discrepancies between yeast complementation assay results and *in planta* function (Lanquar *et al.*, 2004; Cailliatte *et al*., 2010; Bassham & Raikhel, 2000), the possibility remains that LjSEN1 could have a different substrate in the *L. japonicus* plant. If this is the case, GmVTL1 must also transport species other than iron. In this context, other VIT family members transport zinc, manganese and cobalt (Lapinskas *et al.*, 1996; Kim *et al*., 2006; Zhang *et al*., 2012; Labarbuta et al., 2017) and these metals are also important for nitrogen fixation (Dilworth *et al*., 1979; O’Hara, 2001; Kaiser *et al*., 2005).

Some characterised VIT proteins transport ferrous rather than ferric iron (Labarbuta et al. 2017; Kato et al 2019). Our assays have not distinguished between these iron species. Isolated soybean symbiosomes can take up both ferrous and ferric iron (Moreau et al., 1995; LeVier et al., 1996; Moreau et al., 1998). If *L. japonicus* has similar systems for iron uptake into symbiosomes, then the fact that mutation of *Sen1* completely abolishes N_2_-fixation (Suganuma *et al*., 2003; Hakoyama et al., 2012) suggests that it is the sole transporter for iron into symbiosomes and further, that both ferrous and ferric iron are transported by LjSEN1 and possibly GmVTL1. This requires further investigation.

Although we established that GmVTL2 can transport iron, we didn’t determine which nodules cells it is expressed in, nor determine if it can complement the *Ljsen1-1* mutant, so we cannot confirm its importance for the symbiosis. However, since EgVIT1 is proposed to function as a dimer (Kato et al. 2019), it is possible that it dimerises with GmVTL1 in planta. It’s expression is half that of GmVTL1, albeit in the same tissues. It appears the two genes arose as the result of duplication of an ancestral gene followed by whole genome duplication. The reduced capacity for iron transport and reduced expression levels may indicate that *GmVTL2* is undergoing non-functionalization (Roulin et al. 2012) but it could also be undergoing neo-functionalization (Roulin et al. 2012) and actually has another function altogether. Further work is required to investigate this possibility.

## Conclusion

We have functionally characterised GmVTL1 and GmVTL2, members of the VIT family in soybean. Both transport iron and are present on the SM in nodules. GmVTL1 most likely catalyses the final plant-directed step in transport of iron, a micronutrient essential for the symbiosis, from the plant to the nitrogen-fixing bacteroids. The ability of *GmVTL1* to restore nitrogen fixation when expressed in the fix^-^ *Ljsen1* mutant, highlights its importance to the symbiosis. The role of GmVTL2 in soybean is less clear and future studies will focus on whether this protein transports a substrate other than iron and whether its role is different to that of GmVTL1.

## Acknowledgements

This research was funded by the Australian Research Council Discovery Project DP150102264 and Industrial Transformation Research HUB IH140100013, and a Grains Research and Development Corporation scholarship to E.B. The authors acknowledge the facilities, and the scientific and technical assistance, of the Australian Microscopy and Microanalysis Research Facility at the Sydney Microscopy and Microanalysis facility, The University of Sydney and the La Trobe University-Comprehensive Proteomics Platform for providing key infrastructure and expertise for this study. We are also grateful to Dr Norio Suganuma, Aichi University of Education for providing the *Lotus japonicus* sen1-1 mutant, Dr Doris Rentsch, University of Bern, for providing the pDR196GW vector and to Catherine Curie, French National Centre for Scientific Research for sharing the updated methods for Perls/DAB staining.

## Author contributions

E.B. contributed to most experimental procedures described and was involved in designing the experiments and writing the manuscript. A.G. was involved in production of transgenic plants and the microscopy for GFP localisation and GUS analysis. F.B. completed the Perls/DAB staining and microscopy to localise iron in *L. japonicus* nodules, isolated microsomal membrane samples, and conducted yeast complementation experiments. M.U., I. T-J and I.K. hosted and assisted E.B. to complete parts of the work on LjSEN1. D.D. contributed to the design of the project, isolated symbiosome membrane samples and wrote parts of the manuscript. P.S. conceived the project, was involved in experimental design and analysis, isolated symbiosome and microsomal membrane samples and wrote parts of the manuscript. All authors contributed to editing of the manuscript.

