## Supporting Information for "GmVTL1 is an iron transporter on the symbiosome membrane of soybean with an important role in nitrogen fixation"

Article acceptance date: Click here to enter a date.

The following Supporting Information is available for this article:

**Table S1** Primer sequences

**Fig. S1** Nitrogenase activity in soybean nodules at 13, 18 and 25 days after inoculation (DAI).

**Fig. S2** GmVTL1 and GmVTL2 transport iron into the vacuole in the BY4741 yeast iron transport mutant *Δccc1*.

**Fig. S3** Localisation of GmVTL1 (a, b) and LjSEN1 (c) in *Saccharomyces cerevisiae Δccc1*.

**Fig. S4** Alignment of LjSEN1, GmVTL1 and GmVTL2 amino acid sequences using ClustalO.

**Fig. S5** Localisation of iron in nodules and infected cells of wild type GIFU and the *sen1* mutant of *L. japonicus.*

**Fig. S6** Comparison of nodules from wild type *L. japonicus* Gifu transformed with empty vector (Gifu-EV), *L. japonicus sen1* transformed with EV (*sen1*-EV) and *L. japonicus sen1* transformed with *G. max VTL1* coding sequence (*sen1-GmVTL1*) expressed from the *G. max* LBc3 promoter.

**Table. S1** Primer sequences. Sequences of primers used in this study to amplify *GmVTL1*, *GmVTL2* and *Ubiquitin* (reference gene) for real time PCR; *GmVTL1*, *GmVTL2*, *AtVIT1*, *LjSEN1* and *MtSEN1* coding sequences (CDS); *GmVTL1*, *GmVTL2* and *LjSEN1* CDS without the stop codon for C-terminal fusion; *GmVTL1* promoter; *AtVIT1* promoter as well as *GmVTL1*, *GmVTL2*, *LjSEN1* and *AtVIT1* CDS for *A. thaliana* complementation.

| **Candidate (amplified)** | **Forward primer** | **Reverse primer** |
|---|---|---|
| GmVTL1 (real time) | TCATTGAAAACTACAGGACTAGGA | CCAAAAGTGATGGACATGG |
| GmVTL2 (real time) | AGAGACTTGGAGACGGAGAAGAG | TAACCCTAGTTCTGTAATCTGCTACGA |
| Ubiquitin (real time) | GTGTAATGTTGGATGTGTTCCC | ACACATTGAGTTCAACACAAACCG |
| GmVTL1 (promoter) | AAAAAGCAGGCTAAAAGAGTGATATCTGTAACTATTAA | GGGGACCACTTTGTACAAGAAAGCTGGGTATGAAGACAAGACTAGAACAA |
| GmVTL1 | GGGGACAAGTTTGTACAAAAAAGCAGGCTCTTGTCTTCATATGGTTGTTGTGTC | GGGGACCACTTTGTACAAGAAAGCTGGGTTTATAATGCGCTAGCACCAAG |

| | | |
|---|---|---|
| (N-terminal CDS) | | |
| GmVTL2 (N-terminal CDS) | GGGGACAAGTTTGTACAAAAAAGCAGGCTTAATGGCTAATGCTACACCTAATGG | GGGGACCACTTTGTACAAGAAAGCTGGGTTTATTATTCTAAAGCGCTAGCACCC |
| GmVTL1 (C-terminal CDS) | GGGGACAAGTTTGTACAAAAAAGCAGGCTCTTGTCTTCATATGGTTGTTGTGTC | GGGGACCACTTTGTACAAGAAAGCTGGGTATAATGCGCTAGCACCAAGTAA |
| GmVTL2 (C-terminal CDS) | GGGGACAAGTTTGTACAAAAAAGCAGGCTTATTTCGCTCTCCCTCCTTTTT | GGGGACCACTTTGTACAAGAAAGCTGGGTATTCTAAAGCGCTAGCACCCATT |
| LjSEN1 (CDS) | GGGGACAAGTTTGTACAAAAAAGCAGGCTTACCACCCTCTTCATCTCTCAAG | GGGGACCACTTTGTACAAGAAAGCTGGGTATTATATTTCCAACCCACCAGCA |
| MtSEN1 (CDS) | AAAAAGCAGGCTCCATGGCTTCTCTTGGTAACCATAAT | AGAAAGCTGGGTATCAAAGTGAACTATAGCCAACGAA |
| AtVIT1 (CDS for yeast) | GGGGACAAGTTTGTACAAAAAAGCAGGCTTAATGTCGTCGGAGGAAGATAAGA | GGGGACCACTTTGTACAAGAAAGCTGGGTACTAATGTTGCACAACTTTAGCC |
| LjSEN1 (CDS no stop codon) | GGGGACAAGTTTGTACAAAAAAGCAGGCTTACCACCCTCTTCATCTCTCAAG | AGAAAGCTGGGTATATTTCCAACCCACCAGCAC |

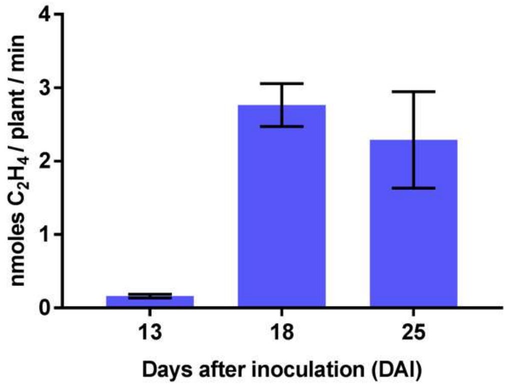


**Fig. S1** Nitrogenase activity in soybean nodules at 13, 18 and 25 days after inoculation (DAI). Acetylene reduction assay was performed on the whole root system of individual plants. The data presented is an average of four biological replicates and error bars represent standard error.

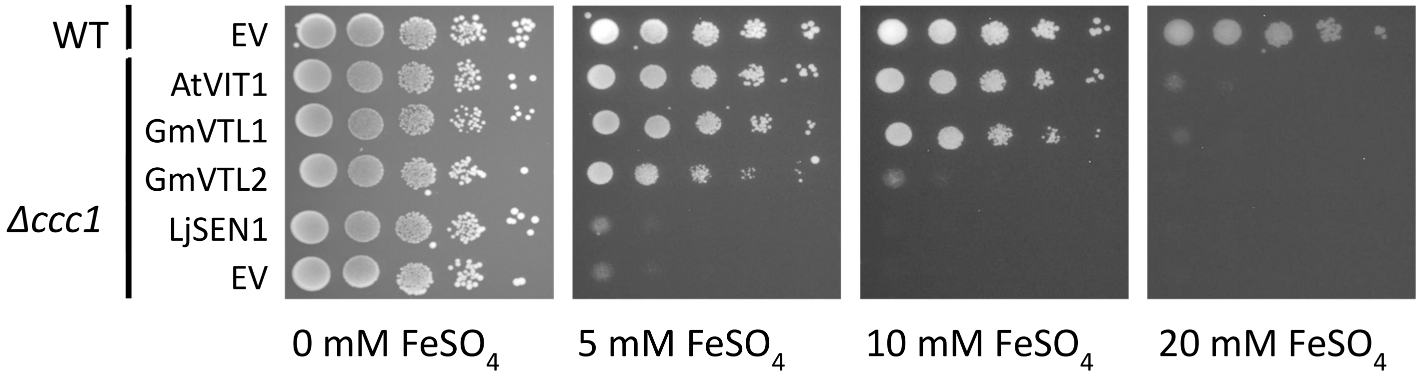


Fig. S2 **GmVTL1 and GmVTL2 transport iron into the vacuole in the BY4741 yeast iron transport mutant *Δccc1*.** Wild type (WT) BY4741 yeast was transformed with the empty vector plasmid (EV; pDR196-GW. The Δccc1 yeast strain derived from BY4741 was transformed with pDR196-GW vector containing the full-length coding sequence of *A. thaliana VIT1* (AtVIT1, positive control), *G. max VTL1 (*GmVTL1), *G. max VTL2* (GmVTL2), and EV, under control of the constitutive PMA promoter. Serial dilutions of each yeast transformant were applied to SD plates (that include 1.6 µM $FeCl_3$) with the addition of 0, 5, 10 or 20 mM $FeSO_4$ and the plates grown for 3 days.

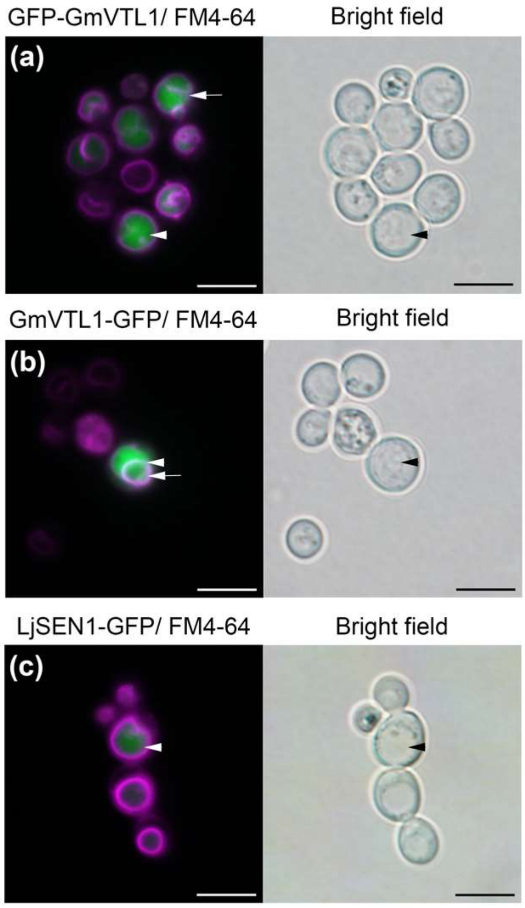


**Fig. S3** Localisation of GmVTL1 (a, b) and LjSEN1 (c) in *Saccharomyces cerevisiae Δccc1*. *Δccc1* yeast expressing GFP-GmVTL1 (a), GmVTL1-GFP (b) and LjSEN1-GFP (c). The yeast tonoplast was stained with FM4-64 and appears pink. The scale bar represents 5 µm. GFP is present inside the vacuolar lumen. The GFP fusion also appears to co-localise with the FM4-64 tonoplast membrane in cells expressing GFP-GmVTL1 and GmVTL1-GFP. LjSEN1-GFP doesn't appear to localise to the tonoplast. The arrowhead indicates the vacuole, the arrow points to regions of co-localisation between GFP and the FM4-64 stained tonoplast.

```
AtVIT1    -------MSSEED------K-I-------TRISIEPEK--QTLLDHHTEKHFTAGEIVRD      37
EgVIT1    -------MADGA-------N-D-------GGNPGAEEQ--QRLLDQHKEAHFTAGEIVRD      36
AtVTL1    ---------------MESHN-V------SNSLN-----LDMEMDQEKAFDYSKRAQWLRA      33
LjSEN1    -------MAAGA------PNHVGTIPNPANG-TTDQNKQTQVTDVPRVVDYWQRAQWLRA      46
GmVTL2    -------MANATPNGSVPHNHVGAVLPTIPTIKIDEKQTLATSEDHTSIDYLQRAQWLRA      53
GmVTL1    MVVVSPKMANATPNGSVPHNHVGAVLLTIPTIKIDGKQTLAT-EDHTSIDYLQRAQWLRA      59
                             :                              :   .: :*

AtVIT1    IIIGVSDGLTVPFALAAGLSGANASSSIVLTAGIAEVAAGAISMGLGGYLAAKSEEDHYA      97
EgVIT1    IIIGVSDGLTVPFALAAGLSGANASSSIVLTAGIAEVAAGAISMGLGGYLAAKSEADNYA      96
AtVTL1    AVLGANDGLVSTASLMMGVGAVKQDVKVMILSGFAGLVAGACSMAIGEFVSVYSQYDIEV      93
LjSEN1    AVLGANDGLVSVASLMMGVGAVNKDAKAMLLAGFAGLVAGACGMAIGEFVSVFTQYEVEV      106
GmVTL2    AVLGANDGLVSVTSLMMGVGAVKKDAKAMLVAGFAGLVAGACGMAIGEFVAVCTQYEVEL      113
GmVTL1    AILGANDGLVSVASLMMGVGAVKRDAKAMLLAGFAGLVAGACGMAIGEFVAVYTQYEVEV      119
           ::*..***.   :*  *:...: . . :: :*:* :.*** .*.:* :::. :: :

AtVIT1    REMKREQEEIVAVPETEAAEVAEILAQYGIEPHEYSPVVNALRKNPQAWLDFMMRFELGL      157
EgVIT1    RELKREQEEIIRVPDTEAAEVAEILARYGIEPHEYGPVVNALRKKPQAWLDFMMKFELGL      156
AtVTL1    AQMKRENGGQ------------------------------------------------VEKE      107
LjSEN1    GQMKRDMIKSE-----------------------------------QGERDLEMAMEKRK      131
GmVTL2    GQMKREMNMSE-----------------------------------GGERDLE--TEK-R      135
GmVTL1    GQMKRDMNMSV-----------------------------------GGERDLEMEMER-R      143
           ::**:

AtVIT1    EKPDPKRALQSAFTIAIAYVLGGFIPLLPYMLIPHAMDAVVASVVITLFALFIFGYAKGH      217
EgVIT1    EKPDPKRALQSAFTIAIAYVLGGLVPLIPYMFIPVARKAVVASVILTLMALLIFGYAKGY      216
AtVTL1    KLP---SPMQAAAASALAFSLGAIVPLMAAAFVKDYHVRIGAIVAAVTLALVMFGWLGAV      164
LjSEN1    SIP---NPMQAALASAFSFSIGGLVPLLSGSFVSVYKIRLLVIVVVVSLALVAFGSVGSL      188
GmVTL2    TLP---NPLQATWASALSFSIGALVPLLSAAFVADYRTRVIVVVAMASLALVVFGSVGAQ      192
GmVTL1    TLP---NPLQATLASALCFSIGALVPLLSAAFIENYRTRIIVVVAMSCLALVVFGWVGAK      200
            *     :*:: : *:.: :*.::**:   ::      : . *    :**. **   .

AtVIT1    FTGSKPLRSAFETAFIGAIASAAAFCLAKVVQH-----    250
EgVIT1    FTDNKPFKSALQTALIGAIASAAAFGMAKAVQS-----    249
AtVTL1    LGKAPVFKSSARVLIGGWLAMAVTFGLTKLIGTHSL--    200
LjSEN1    LGKTPMTRSCVRFMIGGWMAMAITFGLTKLLSAGGLEI    226
GmVTL2    LGKTPKLKSCVRFLLGGWIAMSITFGLTKLMGASALE-    229
GmVTL1    LGKTPKLKSCVRFLLGGCIAMSITFGSTKLLGASAL--    236
          :      :*. .  : * :* : :*  :* :
```

Fig. S4 Alignment of LjSEN1, GmVTL1, GmVTL2, AtVTL1, AtVIT1 and EgVIT1 amino acid sequences using ClustalO. “*” indicates an amino acid conserved in all sequences; “:” indicates conservation between groups of amino acids with strongly similar properties; “.” indicates conservation between groups of amino acids with weakly similar properties. A54, R124 and G202 in GmVTL1, which are conserved in LjSEN1 and when substituted interfere with iron transport in yeast are indicated in orange. E32 in EgVIT1 is highlighted in green and is important for ion translocation from the metal binding domain to the transmembrane domain. E102 and M149 in EgVIT1 are highlighted in red and are involved in ion binding in the metal binding domain (Kato et al. 2019).

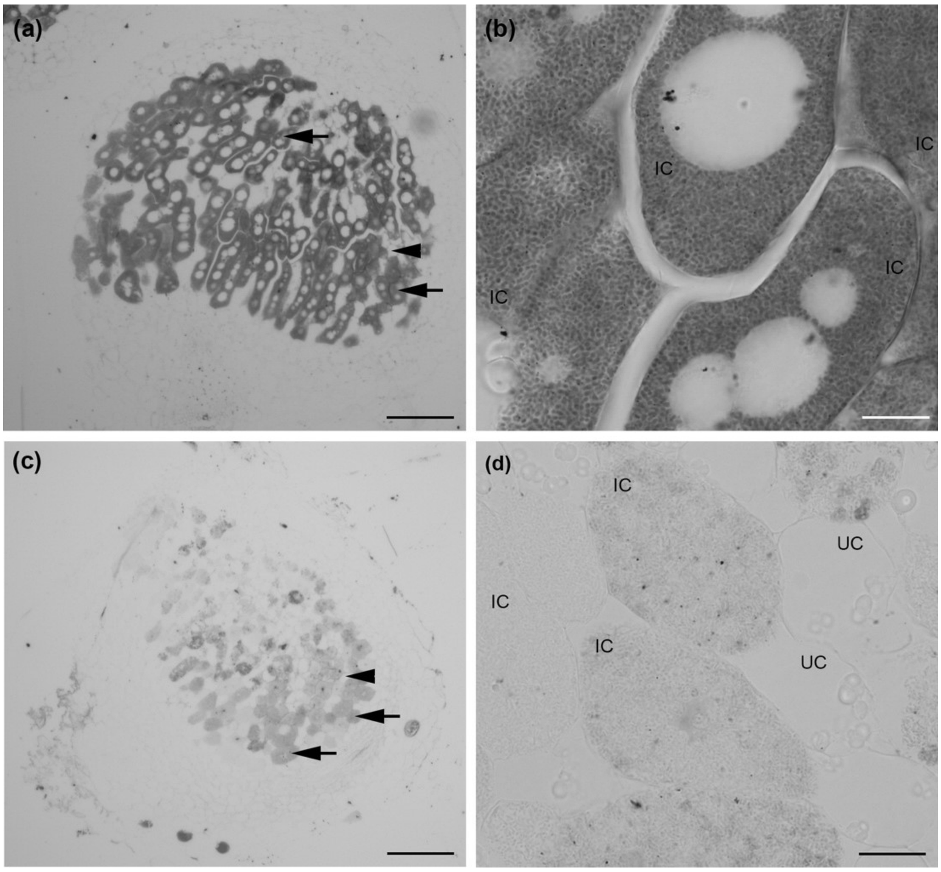


**Fig. S5** Localisation of iron in nodules of wild type *L. japonicus* GIFU (a and b) and the *sen1* mutant (c and d). Sections were stained with Perls/DAB. Arrows indicate infected cells (IC). Arrow heads indicate uninfected cells (UC). Scale bars are 100 µm (a and c) and 10 µm (b and d).

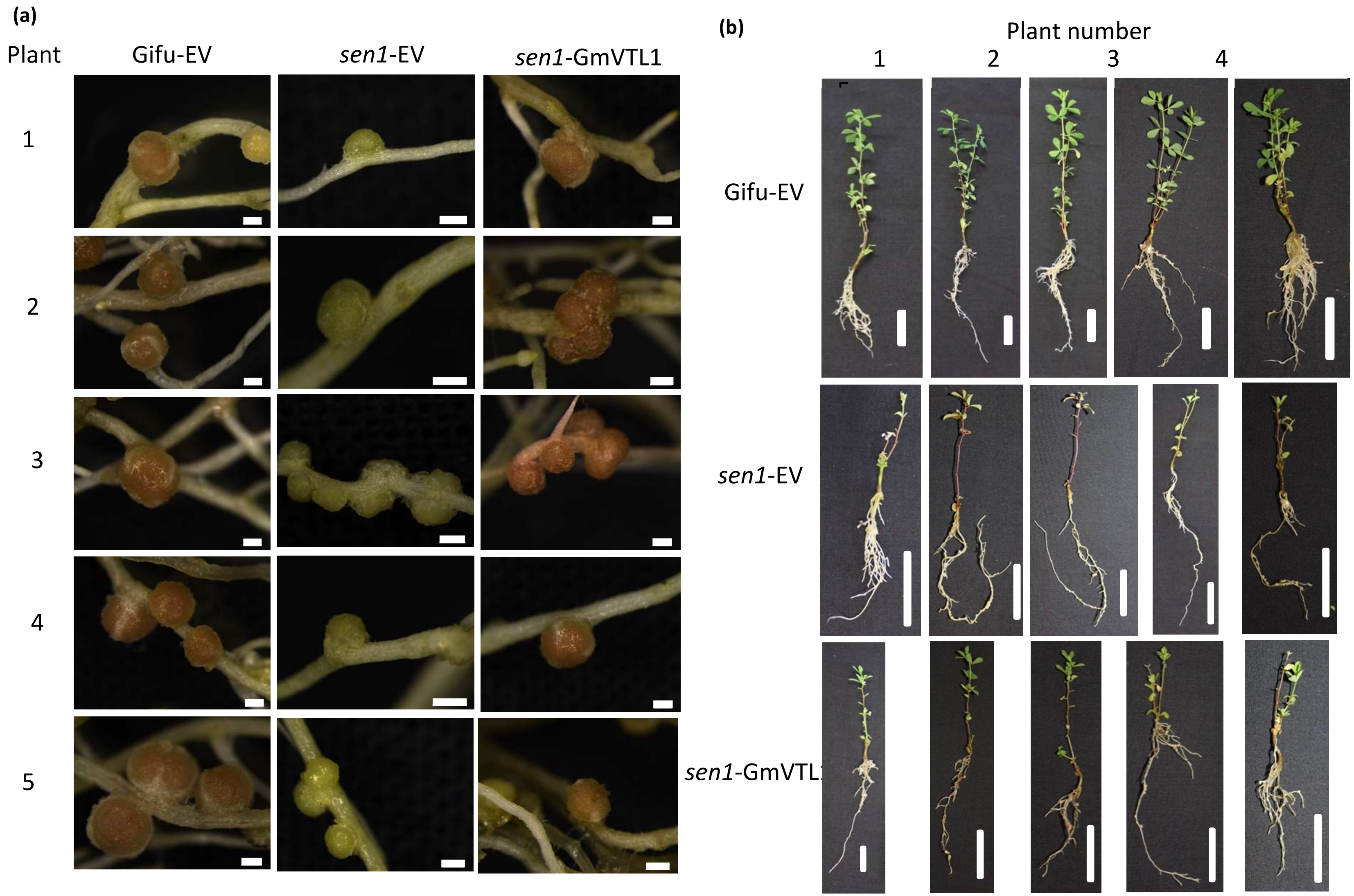


**Fig. S6** Comparison of nodules from wild type *L. japonicus* Gifu transformed with empty vector (Gifu-EV), *L. japonicus sen1* transformed with EV (*sen1*-EV) and *L. japonicus sen1* transformed with *G. max VIT1* coding sequence (*sen1-GmVTL1*) expressed from the *G. max* LBc3 promoter. Images of five complemented plants **(a)** nodules (scale bars represent 500 µm) and (b) whole plants (scale bars represent 2 cm).
